# Mfd regulates RNA polymerase association with hard-to-transcribe regions *in vivo*, especially those with structured RNAs

**DOI:** 10.1101/2020.05.28.121731

**Authors:** Mark N. Ragheb, Christopher Merrikh, Kaitlyn Browning, Houra Merrikh

**Affiliations:** Molecular and Cellular Biology Graduate Program and Medical Scientist Training Program, University of Washington, Seattle, WA, USA; Department of Biochemistry, Vanderbilt University, Nashville, TN 37205, USA

**Keywords:** Mfd, RNA polymerase, regulatory RNAs, RNA secondary structure, toxin-antitoxins

## Abstract

RNA polymerase (RNAP) encounters various roadblocks during transcription. These obstacles can impede RNAP movement, influence transcription, ultimately necessitating the activity of RNAP associated factors. One such factor is the bacterial protein Mfd; a highly conserved DNA translocase and evolvability factor that interacts with RNAP. Although Mfd is thought to function primarily in the repair of DNA lesions that stall RNAP, increasing evidence suggests that it may also be important for transcription regulation. However, this is yet to be fully characterized.

To shed light on Mfd’s *in vivo* functions, we identified the chromosomal regions where it associates. We analyzed Mfd’s impact on RNAP association and transcription regulation genome-wide. We found that Mfd represses RNAP association at many chromosomal regions. We found that these regions show increased RNAP pausing, suggesting that they are hard-to-transcribe. Interestingly, we noticed that the majority of the regions where Mfd regulates transcription contain highly structured regulatory RNAs. The RNAs identified regulate a myriad of biological processes, ranging from metabolism, to tRNA regulation, to toxin-antitoxin (TA) functions. We found that transcription regulation by Mfd, at least at some TA loci, is critical for cell survival. Lastly, we found that Mfd promotes mutagenesis in at least one toxin gene, suggesting that its function in regulating transcription may promote evolution of certain TA systems, and other regions containing strong RNA secondary structures. We conclude that Mfd is an RNAP co-factor that is important, and at times critical, for transcription regulation at hard-to-transcribe regions, especially those that express structured regulatory RNAs.

**Significance:** The bacterial DNA translocase Mfd binds to stalled RNAPs and is generally thought to facilitate transcription-coupled DNA repair. Most of our knowledge about Mfd is based on data from biochemical studies. However, little is known about Mfd’s function in living cells, especially in the absence of exogenous DNA damage. Here, we show that Mfd modulates RNAP association and alters transcription at a variety of chromosomal loci, especially those containing highly structured, regulatory RNAs. As such, this work improves our understanding of Mfd’s function in living cells, and assigns it a new function as a transcription regulator.

## Introduction

Timely and efficient transcription is a fundamental requirement for maintaining cellular homeostasis. Previous work has shown that transcription elongation is discontinuous, with RNA polymerase (RNAP) processivity being altered by a wide range of obstacles(1, 2). These impediments vary in severity, from pause sites that slow the rate of RNAP movement(3–5) to more severe obstacles, such as protein roadblocks and the replication fork. These impediments can induce reverse translocation of RNAP with respect to both DNA and the nascent RNA (RNAP backtracking)(6–9). The impact of these roadblocks on RNAP processivity are prevented and resolved through various mechanisms including the coupling of transcription and translation, as well as various cellular factors that help re-establish transcription elongation (i.e. anti-backtracking factors such as GreA)(10).

*In vitro* work shows that the DNA translocase Mfd utilizes its RNAP binding properties and forward translocase activity to rescue arrested RNAPs, restoring transcription elongation(11) as well as promoting transcription termination(12). However, despite decades of research on the biochemical characteristics of Mfd, the endogenous contexts in which its translocase and anti-backtracking functions are critical for transcription remain elusive.

Mfd was initially described as a critical DNA repair factor *in vivo* that promotes transcription-coupled repair (TCR)(13–15). In the TCR pathway, Mfd removes stalled RNAPs at bulky DNA lesions and promotes nucleotide excision repair (NER) via its UvrA binding capacity (see review (16)). However, cells lacking Mfd show little to no sensitivity to DNA damaging agents that promote RNAP stalling(17, 18), especially relative to NER proteins(19, 20). Together with its high degree of conservation, such data suggests that Mfd may have a broader cellular function outside of DNA repair.

Mfd was shown to have other roles in the cell, such as regulating catabolite repression in *Bacillus subtilis*(21, 22). Recent *in vitro* experiments have shown that Mfd is capable of autonomously translocating on DNA in the absence of a lesion, but whether this occurs *in vivo* is unclear(23). Interestingly, Mfd functions as an evolvability factor, promoting mutagenesis and the rapid evolution of antibiotic resistance in diverse bacterial species(17). Mfd also promotes mutagenesis during stationary-phase(24, 25) and at chromosomal regions where collisions between the replication and transcription machineries occur(10). However, a comprehensive *in vivo* study examining what genomic “hotspots” may be prone to Mfd’s mutagenic activity has not been performed.

Despite our limited understanding of Mfd’s cellular functions, its high level of conservation in bacteria implies a fundamental role for Mfd that may be separate from TCR. In addition, Mfd is constitutively expressed, suggesting that it may have homeostatic role in transcription regulation. Further work has recently suggested that Mfd plays a housekeeping function in cells by associating with RNAP in the absence of exogenous stressors(26). However, the field still lacks a clear understanding of the conditions and chromosomal features that prompt Mfd binding and modulation of RNAP association and function.

In this work, we identify the genomic regions of Mfd association in both *B. subtilis* and *Escherichia coli*. We also identify the regions where Mfd modulates RNAP association and transcription. We find that Mfd alters RNAP density predominantly at regions containing highly structured RNAs in both *B. subtilis* and *E. coli*. Our analysis of RNAP pausing experiments strongly suggests that these sites are hard-to-transcribe. We find that Mfd regulates the transcription of genes involved in various cellular functions including toxin-antitoxin (TA) systems. Importantly, we observe that Mfd’s regulatory activity, at least for TA systems, can be essential: when we overexpress the toxin genes, cell viability is significantly compromised without Mfd. Lastly, we find that Mfd promotes mutagenesis of at least the *txpA* toxin gene. We conclude that RNA secondary structure is a major impediment to transcription *in vivo* and that Mfd is an important RNAP co-factor that regulates transcription and RNAP association with such loci.

## Results

### Mapping the genomic loci where Mfd associates

We began by identifying the chromosomal regions where Mfd associates using chromatin immunoprecipitation followed by high-throughput sequencing (ChIP-seq). We constructed a *B. subtilis* strain where Mfd is C-terminally Myc-tagged (Fig. S1). Prior work has shown that C-terminally tagged Mfd retains its functionality *in vivo*(26) – (below we present data that confirms the functionality of this Mfd-Myc in our system). To identify the chromosomal regions where Mfd associates, we harvested exponentially growing cells and performed ChIP-seq analysis. We controlled for potential ChIP artifacts by comparing this signal to ChIP-seq performed using Myc antibody against *B. subtilis* cells lacking the Myc-tagged Mfd (Fig. 1*A*). Under these conditions, 489 out of 5755 genes analyzed in the *B. subtilis* genome exhibited preferential Mfd association (defined as one standard deviation greater than the average Mfd ChIP signal across all genes) (Dataset S1).

**Fig. 1.**
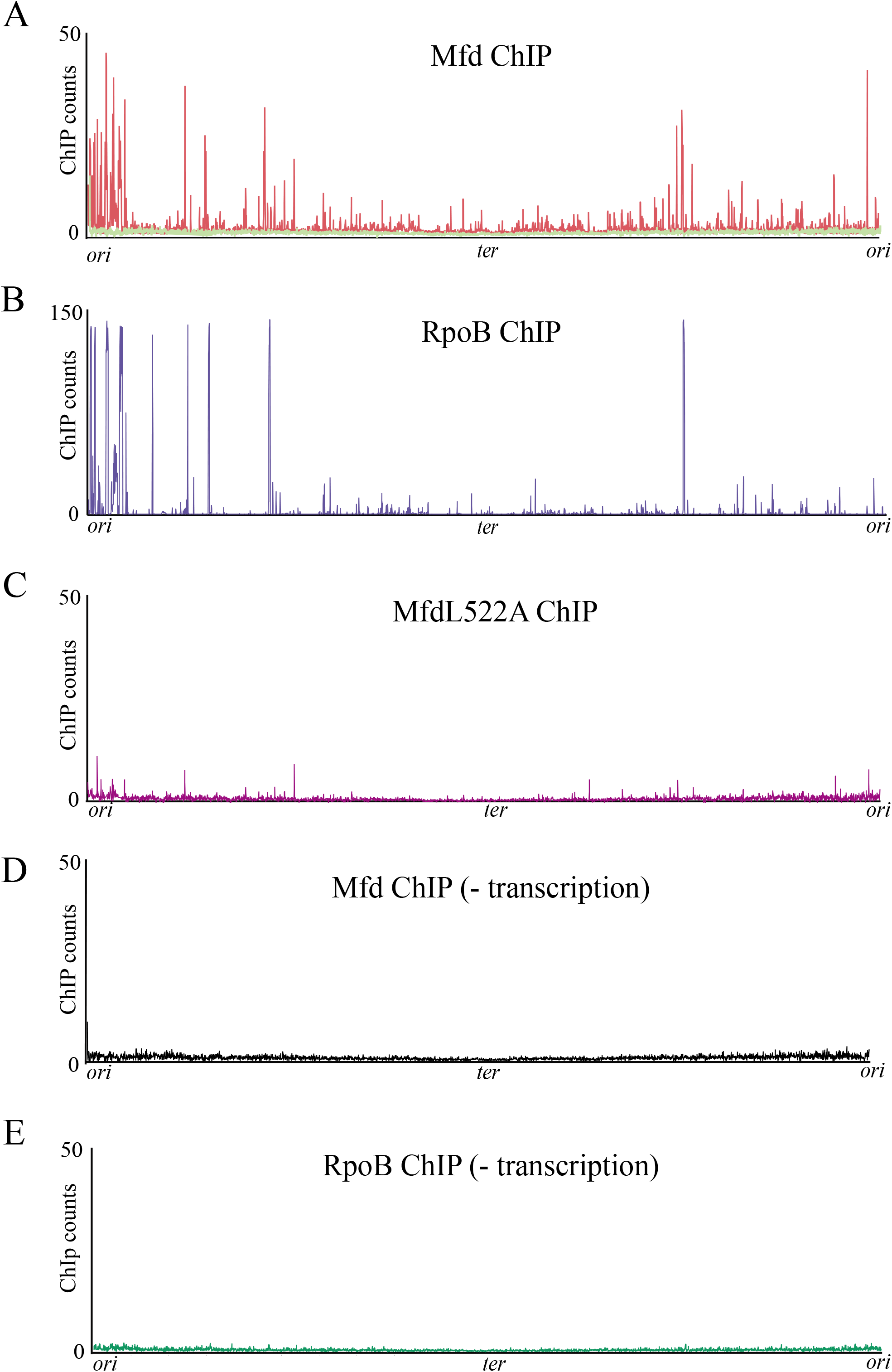
Mfd functions as an RNAP co-factor and requires transcription elongation for association with DNA. (A) ChIP-seq plot of *B. subtilis* Mfd tagged with 1x myc (red) and of WT *B. subtilis* (light green) using myc antibody. (B) ChIP-seq plot of WT *B. subtilis* RpoB. (C) ChIP-seq plot of *B. subtilis* MfdL522A-myc point mutant. (D) *B. subtilis* Mfd-myc and (E) *B. subtilis* RpoB ChIP-seq after treatment with 50μg/mL of rifampicin for five minutes. Plots are normalized to total DNA input controls and are the average of at least two independent experiments.

### Mfd’s genomic association pattern correlates with that of RNAP

Given the known physical interaction between Mfd and RNAP(27), we hypothesized that Mfd binding sites may correspond to RNAP binding locations. To test this, we performed ChIP-seq of RpoB, the β subunit of RNAP, using a native antibody. Indeed, we found that Mfd association largely overlaps with RpoB occupancy (Pearson coefficient = 0.68) (Fig. 1*B* and Fig. S2), consistent with Mfd’s proposed function as an RNAP co-factor in *B. subtilis*.

### Mfd requires interaction with RNAP for its association with all genomic loci

Mfd is a multi-modular protein, consisting of eight domains connected by flexible linkers(27). Of these domains, the RNAP interacting domain (RID) and the translocase module (composed of domains D5 and D6) are critical for Mfd’s ability to rescue stalled transcription complexes(28–30). *In vitro*, Mfd is recruited to the identified genomic regions through its interaction with RpoB. We therefore tested whether the interaction of Mfd with RpoB is critical for its recruitment to the genomic loci we identified. Prior *in vitro* work suggested that RID mutations abrogate the interaction between Mfd and RNAP(27). We therefore constructed a strain containing a Myc tagged Mfd, with a mutation at the L522 residue to disrupt Mfd’s binding to RpoB, without disrupting the stability or folding of Mfd, analogous to that described in *E. coli*(27). Upon confirming that the *B. subtilis* L522A mutation (analogous to L499R mutation described in *E. coli*) disrupted Mfd’s interaction with RNAP via a bacterial 2-hybrid assay (Fig. S3), we performed ChIP-seq experiments with this mutant. The ChIP signal we detected in WT strains was abrogated in the strain expressing the L522A allele of *mfd* (Fig. 1*C*). These results strongly suggested that Mfd’s interaction with RNAP is essential for its recruitment to all genomic loci identified in the previous experiments and that Mfd functions as a genome-wide RNAP co-factor *in vivo*.

### Mfd’s association with DNA requires transcription elongation

*In vitro*, Mfd helps promote the rescue of arrested transcription elongation complexes (TECs), yet how Mfd recognizes stalled RNAPs *in vivo* remains unclear. We therefore asked whether Mfd association with various genomic loci was facilitated via loading during the transcription initiation or elongation phase. To distinguish between transcription initiation and elongation, we utilized the antimicrobial rifampicin, which directly blocks transcription initiation(31, 32), subsequently eliminating the formation of transcription elongation complexes (TECs). We performed both Mfd and RpoB ChIP-seq studies in the presence of rifampicin and found that this treatment largely eliminated both Mfd and RpoB ChIP-seq binding signal (Fig. 1*D-E*). This finding is consistent with biochemical evidence showing that Mfd is unable to release RNAP at initiation sites(11). These data suggest that Mfd associates with elongating rather than initiating RNAPs *in vivo*.

### Mfd decreases RNAP density at some genomic loci

Mfd can function to promote both transcription elongation as well as transcription termination, at least *in vitro* (11, 12). However, the importance of these functions *in vivo* remains elusive. Because of Mfd’s effects on RNAP *in vitro*, we wondered if and how Mfd’s close association with RNAP occupancy *in vivo* altered RNAP association in living cells. We therefore performed ChIP-seq of RpoB in WT and *Δmfd* strains and identified where RpoB occupancy is altered in the absence of Mfd. ChIP-seq experiments did not detect alterations in RpoB occupancy at the majority of genes where Mfd associates based on our Mfd ChIP-seq studies. This may be a result of various factors, such as the existence of redundant transcription-associated factors, the specific growth conditions of the experiment, or potential limitations of detection thresholds which are possible in an ensemble assay such as ChIP-seq. However, we did find a number of genes where RpoB occupancy either increased or decreased in the *Δmfd* strain compared to WT (Fig. 2*A*). We specifically noticed a bias towards a greater number of genes that exhibited increased rather than decreased RpoB occupancy in the *Δmfd* strain. Quantification of these results revealed that a total of 116 genes exhibited at least a two-fold increase, while 53 genes exhibited at least a two-fold decrease in RpoB occupancy without Mfd. Many of these genes are within the same operon, and therefore are expressed as single transcripts. Thus, rather than single genes, we grouped and analyzed the identified genes as transcription units (TUs)(33).

**Fig. 2.**
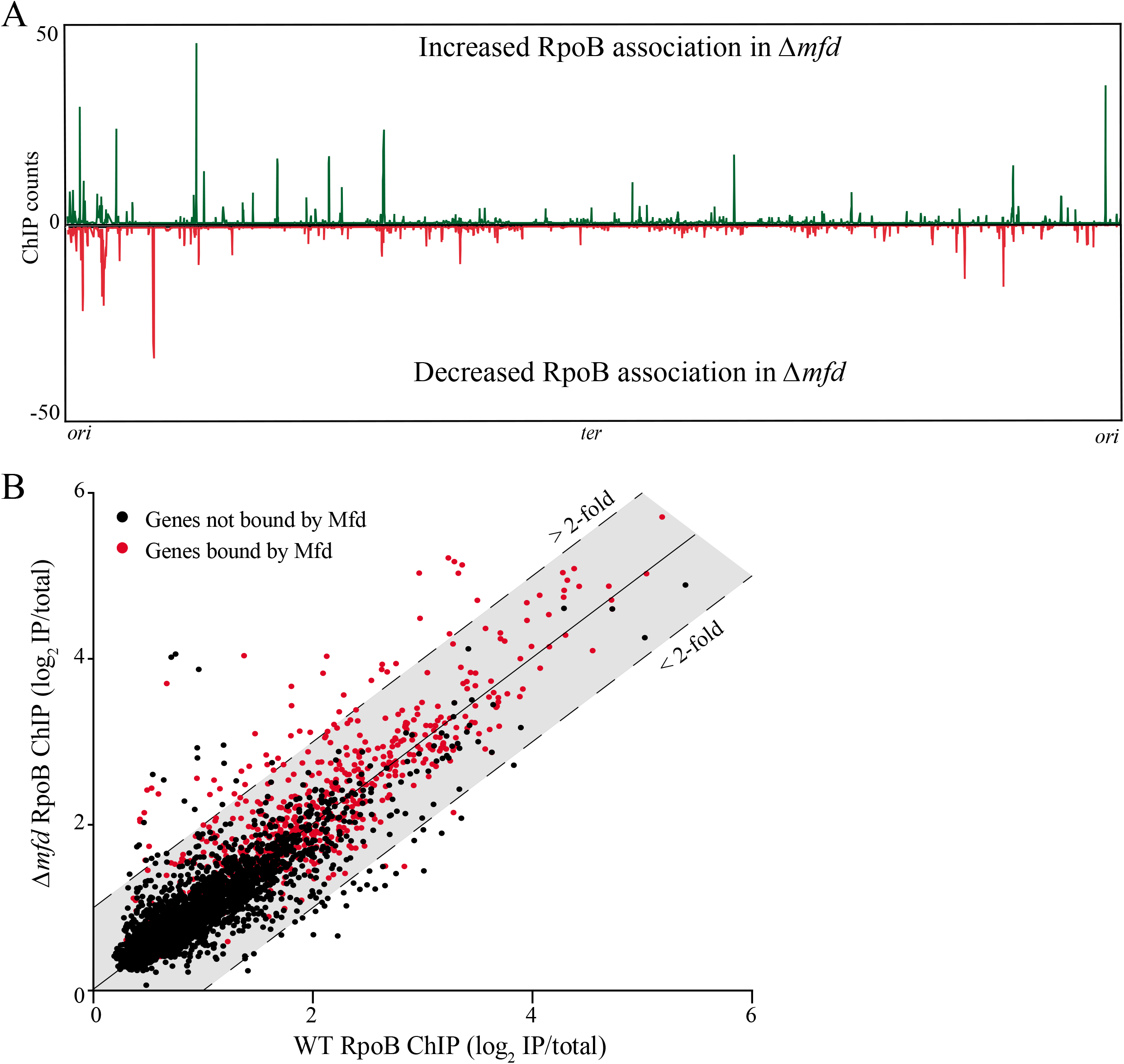
Mfd directly promotes release of RNAP *in vivo*. (A) RpoB ChIP-seq plots showing regions of RpoB enrichment in *Δmfd*. Top half of graph (counts in green) reflects normalized RpoB ChIP-seq read counts where *B. subtilis Δmfd* had increased signal relative to WT. Bottom half of graph (counts in red) reflects RpoB ChIP-seq read counts where *Δmfd* had decreased signal relative to WT *B. subtilis*. High background signal from ribosomal RNA was removed from plots for better visualization. (B) Scatter plot of WT and *Δmfd* RpoB ChIP-seq measuring signal at each gene in *B. subtilis*. For quantification of ChIP signal at each gene, read counts for each gene were normalized to total library counts and IP samples were normalized to total DNA input to calculate an IP/Total DNA ratio. Ratios were log2 normalized and averaged over at least two independent experiments. Data points above and below colored shading indicate greater than two-fold increase and decrease in RpoB signal in the *Δmfd* strain, respectively. Data points in red indicate genes bound by Mfd. Binding is defined as one standard deviation greater than the average ChIP signal across all genes in *B. subtilis*. Calculation of Mfd binding at each gene was determined as described for RpoB ChIP samples.

Our analysis revealed that *Δmfd* strains contain 71 TUs with at least one gene containing a minimum of two-fold increase, and 31 TUs with at least one gene containing a minimum of two-fold decrease in RpoB occupancy compared to WT (Table 1, Datasets S2 and S3).

**Table 1.** Summary list of genes and previously defined TUs bound with changes in RpoB density in *Δmfd* from ChIP-seq analysis. Changes in RpoB density and Mfd binding at TUs are defined by changes in one or more genes corresponding to its associated TUs

|  | Increased RpoB density in<br>*Δmfd* | Decreased RpoB density in<br>*Δmfd* |
| --- | --- | --- |
| **Genes (and TUs)** | 116 (71) | 53 (31) |
| **Genes (and TUs) bound**<br>**by Mfd** | 52 (35) | 0 (0) |

We next wanted to determine whether the changes in RpoB occupancy observed in the *Δmfd* strain were directly due to Mfd’s activity at those regions or whether we were detecting indirect effects. To address this, we looked for a correlation between the regions where Mfd associates (ChIP-seq experiments) and where there are also changes in RpoB association levels in the absence of Mfd. We only observed Mfd association at the genes that showed increased RpoB occupancy in the *Δmfd* strain. The genes with decreased RpoB occupancy in the *Δmfd* strain did not show Mfd association (Fig. 2*B*). Specifically, we detected Mfd at 52 of the 116 genes with increased RpoB association (35 of the 71 TUs) in the *Δmfd* strain compared to WT. We did not detect Mfd association at any of the 53 genes (31 TUs) with decreased RpoB occupancy. Because these 31 TUs do not display Mfd association, we concluded that decreased RpoB occupancy at these sites are not directly due to Mfd activity and likely reflect indirect effects on transcription in the *Δmfd* strains.

We next performed RpoB ChIP-qPCR analysis to confirm our ChIP-seq results. We chose two sites from our candidate genes with Mfd binding and increased RpoB occupancy in the *Δmfd* strain. We found results consistent with our ChIP-seq studies (Fig. S4). We additionally tested the functionality of the Myc tagged Mfd using RpoB occupancy as our readout. We did not find changes in the RpoB levels in the tagged Mfd strain, confirming its functionality, at least in regulating RNAP association (Fig. S4).

Based on these findings, we conclude that increased RNAP occupancy at the identified chromosomal sites is a direct result of Mfd’s function as either an RNAP termination (12) or processivity factor (34). In other words, the data strongly suggest that under regular circumstances, Mfd decreases RNAP association.

### Regions where Mfd increased RNAP density are enriched for regulatory RNAs

*In vitro*, Mfd’s translocase activity can help release RNAP or assist it with elongation when exposed to different obstacles. However, whether there are endogenous hotspots of RNAP stalling that require Mfd function remains unknown. Furthermore, if such hotspots exist, the nature of the potential obstacles remains to be determined. Intriguingly, 92% of the TUs that showed both an increase in RNAP density in the *Δmfd* strain and direct Mfd association express a minimum of one regulatory RNA (Table S1). These regulatory RNAs are a subset of the 1583 regulatory RNAs in *B. subtilis*, which encompass a wide variety of RNAs, including independent, non-coding transcripts, antisense RNAs, and multiple riboswitches(33). In comparison, only 39% of the TUs with decreased RNAP density in *Δmfd* strains contain regulatory RNAs (Table S2), which is consistent with the average percentage of TUs with predicted regulatory RNAs in *B. subtilis*(33, 35).

### Sites of Mfd function contain highly structured regulatory RNAs

We hypothesized that Mfd function at the identified regions was related to RNA secondary structure impeding RNAP processivity. This hypothesis is consistent with changes in RNAP processivity due to secondary RNA structures, such as hairpins in the context of intrinsic transcription termination, and other transcription regulatory processes(36–38). Previous work characterized the predicted secondary structure for each regulatory RNA in *B. subtilis*(35). We sought to test whether the regions with increased RNAP in *Δmfd* strains are more prone to transcribing RNAs with more stable secondary structures. We determined the average minimum free energy (MFE) z-score for each of the RNAs at these regions as a proxy of RNA structure stability(39, 40). Specifically, we examined the regulatory RNAs in the TUs that had both increased RpoB density in the *Δmfd* strain and Mfd association. We then compared the results to TUs that showed no difference in RpoB density between WT and *Δmfd* strains. We found that the TUs where there is Mfd binding and increased RpoB density in the *Δmfd* strain have significantly higher predicted RNA secondary structures relative to all other known regulatory RNAs (Fig. 3*A*).

**Fig. 3.**
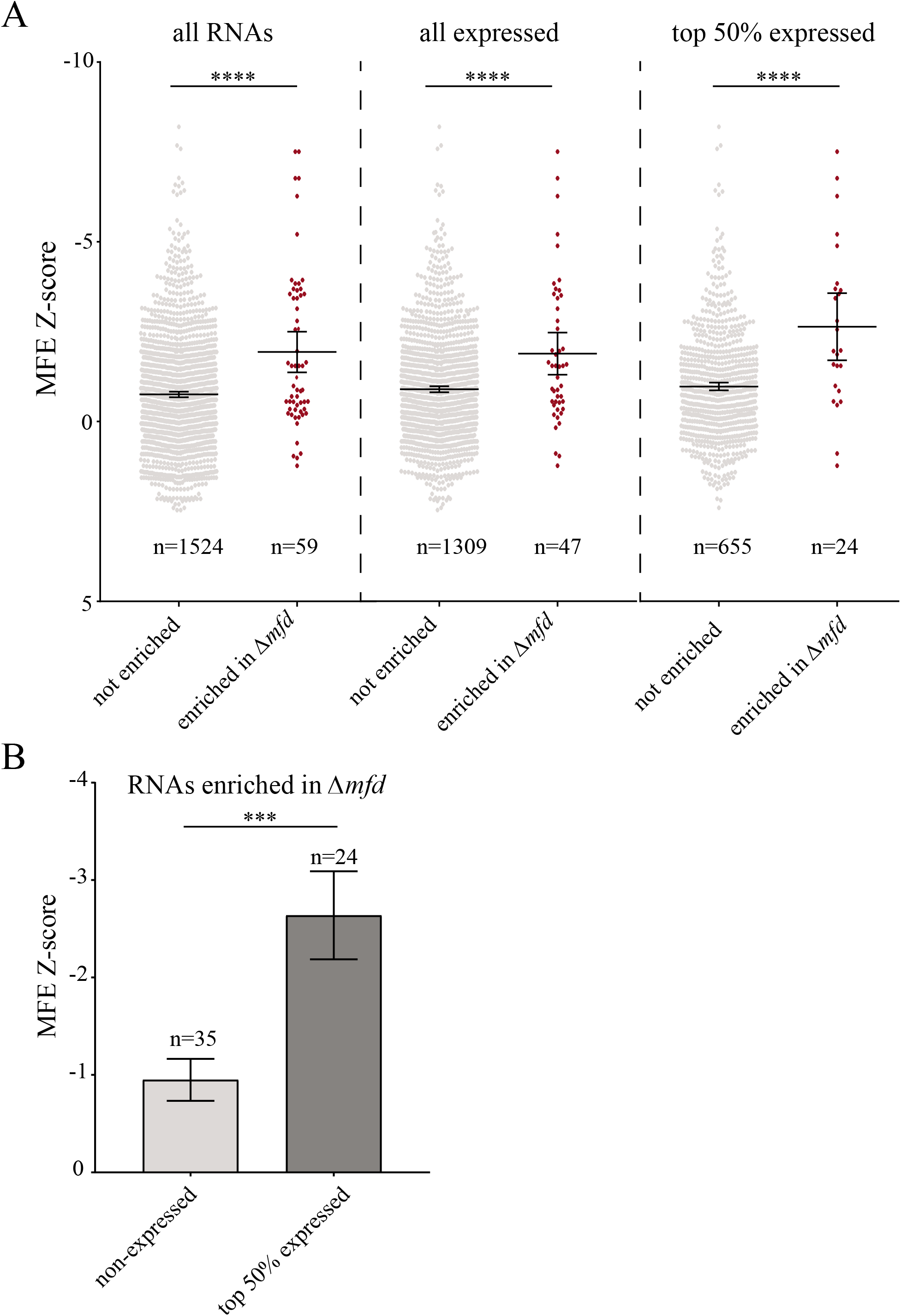
Transcription units with Mfd binding and increased RNAP density in *Δmfd* are enriched for structured regulatory RNAs. (A) Scatter plot of the minimum free energy (MFE) Z-score for regulatory RNAs in *B. subtilis*. Data points represent regulatory RNAs within TUs that have no observed change in RpoB density between WT and *Δmfd* (grey points) and TUs that have increased RpoB density in *Δmfd* and are also bound by Mfd (red points). The three scatter plots represent all regulatory RNAs (left), only expressed regulatory RNAs (middle) and top 50% expressed regulatory RNAs (right). Expression data determined from RNA-seq analysis. (B) Bar graph of regulatory RNAs within TUs that have increased RpoB density in *Δmfd* and are bound by Mfd. Non-expressed RNAs shown in light grey and top 50% of expressed RNAs shown in dark grey. Error bars represent the standard error of the mean (SEM). Statistical significance was determined using two-tailed Z-test for two population means (****p<0.0001).

Many of these regulatory RNAs are not transcribed during standard growth conditions. To control for the potential biases of including non-expressed RNAs, we stratified our analysis to exclude RNAs that were not expressed under our growth conditions, as determined from our RNA-seq data (see Fig. S8). Consistent with our global analysis, we find that expressed regulatory RNAs associated with Mfd binding and increased RpoB density in *Δmfd* have significantly higher MFE z-scores (Fig. 3*A*, Fig. S5). We also find that within regulatory RNAs associated with Mfd binding and increased RpoB density in the *Δmfd* strains, those that were highly expressed (top 50%) have a higher MFE z-score compared to those that were not expressed (Fig. 3*B*). These findings suggest that Mfd regulates RNAP at regions containing highly structured RNAs.

### Sites of Mfd activity correlate with difficult-to-transcribe regions

If sites of Mfd association and Mfd alteration in RpoB density were related to RNAP processivity, we would expect these sites to be difficult-to-transcribe. To identify such regions, we utilized experimental data from native elongating transcript sequencing (NET-seq) in *B. subtilis*(41). Through the capture of nascent transcripts from tagged immunoprecipitation of RNAP molecules, NET-seq allows for identification of RNAP pausing in living cells with high resolution(41). We observed that many genes with increased RpoB levels without Mfd displayed high NET-sig signal, and that the corresponding sequencing patterns closely aligned (Fig. 4*A*). We quantified this pattern by analyzing the NET-seq read count across the previously identified genes where Mfd associates and has increased RpoB signal either with or without Mfd. We found that the median NET-seq signal is significantly higher, by ~3.7-fold, at genes with both increased RpoB signal in *Δmfd* strains and Mfd binding relative to genes with decreased RpoB signal (Fig. 4*B-C*).

**Fig. 4.**
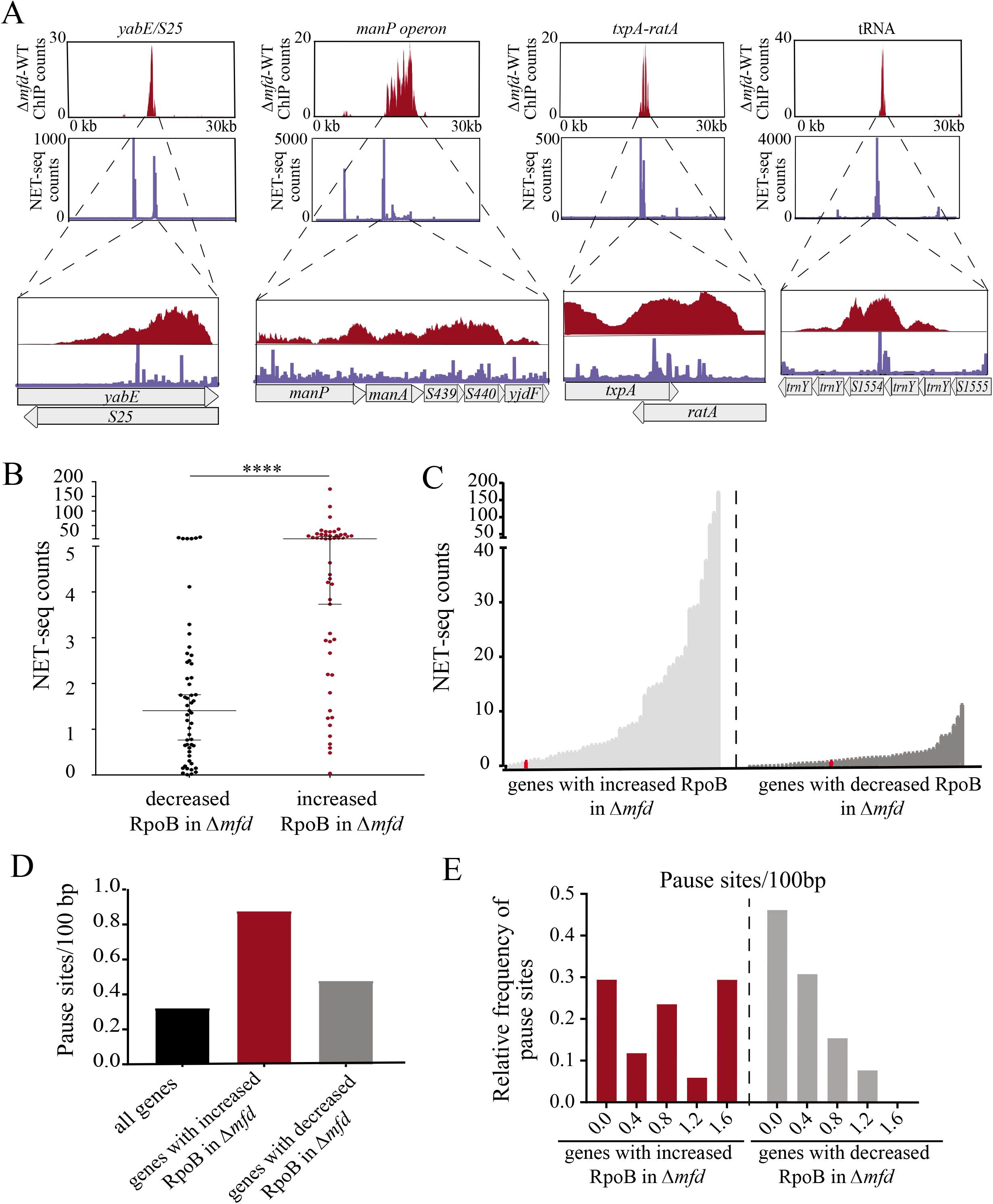
Genes with Mfd binding and increased RNAP density in *Δmfd* are enriched for RNAP pause sites. (A) Read counts for RpoB ChIP-seq (red) and NET-seq (blue) across a 30kb window for four representative genomic regions (*yabE/S25, manP* operon, *txpA/ratA*, and a tRNA locus). Zoomed in plots showing the gene/operon only (y-axis scale is preserved). (B) Scatter plot of the average NET-set read counts for genes with decreased RpoB association in *Δmfd* (black dots, n=53) and increased RpoB association in *Δmfd* (red dots, n=52). Median and 95% confidence interval shown. Statistical significance was determined using the nonparametric Mann-Whitney test for two population medians (**** p<0.0001). (C) Line plot showing by genes with increased (light grey) and decreased (dark grey) RpoB association in *Δmfd*, ranked by increasing signal. Red data point in both plots represents the mean NET-seq read count across the genome. (D) Average number of pause sites (per 100bp) for all genes in *B. subtilis* (black bar), and genes with increased (red bar) and decreased RpoB association (grey bar) in *Δmfd*, respectively. (E) histogram showing the relative frequency of pause sites (per 100bp) in genes with increased (red bars) and decreased (grey bars) RpoB association in *Δmfd*.

Larson et al. identified 9,989 discrete pause sites in *B. subtilis*, defined as nucleotides with greater than four standard deviations in the NET-seq read count(41). We therefore tested whether sites with proposed Mfd activity due to RNA secondary structure contained a greater number of pause sites relative to the average number across the genome. We indeed found that there are roughly 3-fold greater number of discrete pause sites in genes with proposed Mfd activity relative to the average number across the genome, with a notable increase in the frequency of genes with more than one pause site per 100bp at these sites (Fig. 4*D-E*). We did not find this pattern in genes with decreased RpoB association in *Δmfd* strains (Fig. 4*D*-*E*). Critically, the discrete pause sites identified by Larson, et al. in *B. subtilis* are limited to coding regions only, and do not include regulatory RNAs, likely underestimating the number of true pause sites requiring Mfd activity. From these findings we conclude that Mfd functions mainly at RNAP pause sites *in vivo*.

### Mfd’s effect on RNAP at sites of structured regulatory RNAs is conserved in *E. coli*

Mfd is highly conserved across all bacterial phyla, and functional homologs exist throughout all domains of life(42–44). Prior work shows that Mfd function is also highly conserved, at least with regards to its mutagenic activity(17). To test whether Mfd’s genome-wide coupling with RNAP was also conserved in gram-negative species, we performed Mfd and RpoB ChIP-seq experiments in *E. coli*. We found that as in *B. subtilis*, Mfd and RpoB occupancy are highly correlated (Pearson coefficient 0.98) in *E. coli* (Fig. S6*A-C*), showing that Mfd’s function as an RNAP co-factor is likely conserved.

We also determined whether altering RpoB occupancy at sites containing structured RNAs was true in *E. coli*. We compared the signal of the ChIP-seq data for RpoB in WT and *Δmfd* strains in *E. coli* and looked for genes where RpoB occupancy was altered in the absence of Mfd. As in *B. subtilis*, we found a higher number of genes that exhibited at least a two-fold increase in RpoB association in the absence of Mfd, compared to genes with decreased RpoB association (105 versus 24 respectively, Fig. S6*D*, Datasets S4 and S5). 49 of the 105 genes with increased RpoB association also had Mfd association (Dataset S4). Of these 49 sites, we found that roughly 40% either contain regulatory RNAs or are thought to contain a structural element. Three of these sites are TA RNAs (*symR, sokA, sokC*)(45, 46). One is a tRNA (*metZ*) and two are small regulatory RNAs (*spf, gadY*), all of which are thought to contain secondary structures(47–49). Seven sites are repeated extragenic/intragenic palindromes (RIP/REP), which contain extensive secondary structure and are thought to facilitate transcription termination(50, 51), and six are ribosomal proteins which contain structured regulatory elements within their 5’ untranslated regions(52). In contrast, no regulatory RNAs, tRNAs, or ribosomal proteins were found in the genes with decreased RpoB association in the *Δmfd* strains (Dataset S5). These findings suggest Mfd’s function at sites of structured regulatory RNAs is largely conserved across species.

### Genome-wide Mfd binding is correlated with RNA secondary structure

While various methods exist for computational predictions of RNA secondary structure, gathering empirical data on RNA structure continues to be technically challenging, particularly in a high-throughput fashion. The recent development of parallel analysis of the mRNA structure (PARS) has allowed for experimental determination of RNA secondary structure in multiple species, including in *E. coli*(53, 54). This method utilizes enzymatic digestion of single and double-stranded RNA with RNA-specific nucleases and couples this with highly selective cDNA generation of digested RNAs and subsequent deep sequencing(53). Del Campo et al. utilized PARS to establish the RNA secondary structure for roughly 2,500 genes in *E. coli*(54). We therefore wanted to determine whether sites experimentally confirmed to have high RNA secondary structure in *E. coli* correlate with regions of Mfd binding. We observed that across genomic sites containing a high PARS score, both the presence of Mfd association and the magnitude of Mfd binding closely aligned with the magnitude of experimentally determined RNA secondary structure (Fig. 5*A*). To further quantify this, we measured the area under the curve (AUC) for both our Mfd ChIP-seq and the PARS data set to assess whether these two signals were correlated. We indeed found a strong positive correlation between the AUC for our Mfd ChIP-seq and the PARS score (r = 0.6) across the *E. coli* genome (Fig. 5*B*). These data suggest that the presence of RNA secondary structure, at least in *E. coli*, at least partially correlates with Mfd genome-wide association and function.

**Fig. 5.**
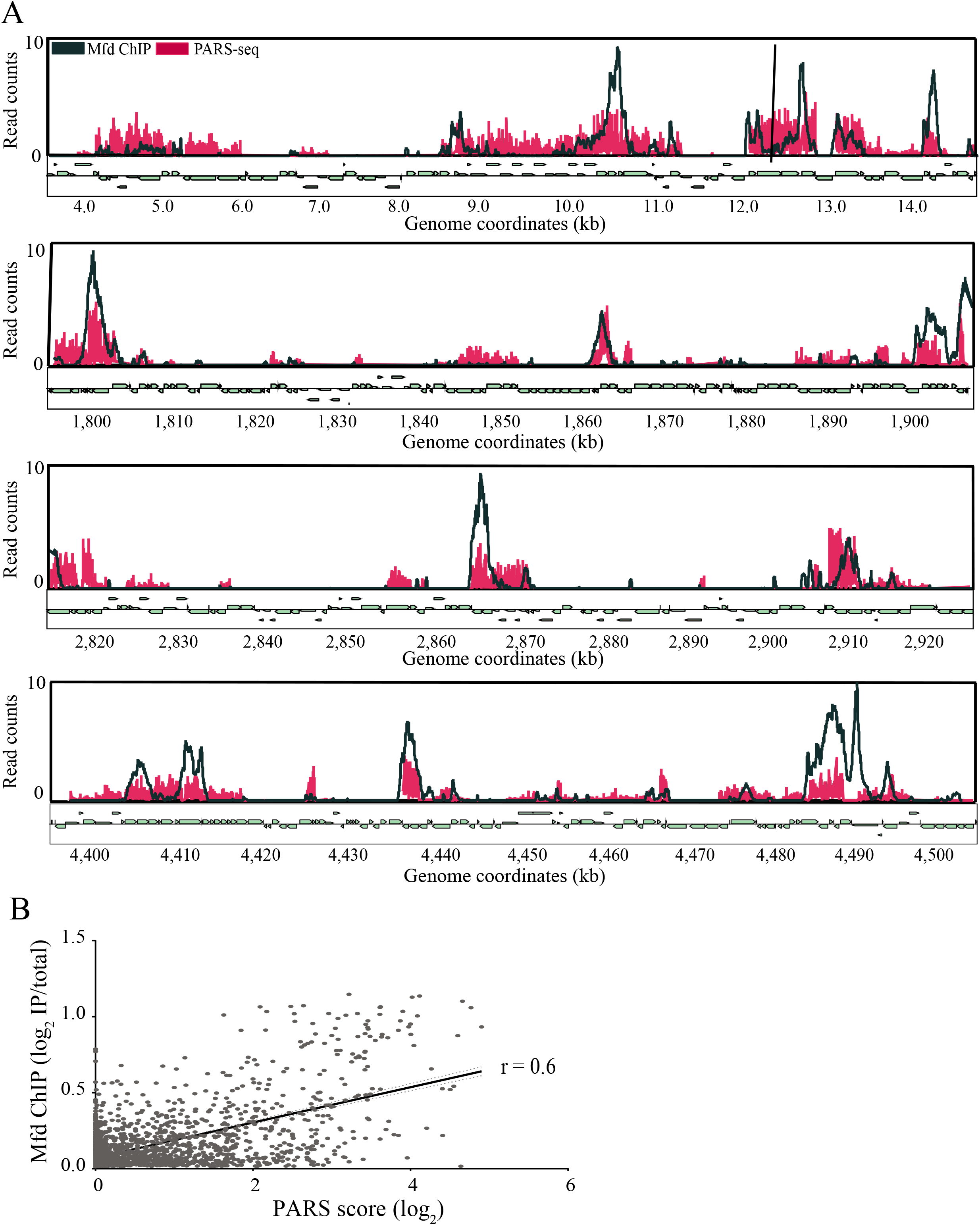
Genome wide Mfd association is correlated with sites of RNA secondary structure based on PARS seq in *E. coli*. (A) Overlay of sequencing counts from Mfd ChIP-seq (dark blue) and PARS-seq (red) across four 100kb regions across the *E. coli* genome. ChIP-seq plots averaged from at least two independent experiments. (B) Linear regression analysis comparing Mfd ChIP-seq and PARS-seq in *E. coli*. Read counts from ChIP-seq and PARS-seq were used to calculate the area under the curve (AUC) for each gene in the *E. coli* genome. Pearson’s correlation coefficient for *E. coli* ChIP-seq and PARS-seq r = 0.6. Dotted lines represent 95% confidence intervals.

### GreA does not preferentially effect transcription of regions containing structured RNAs

Various factors are known to help rescue arrested RNAP through different mechanisms. One of the most well-known and highly conserved proteins is GreA, which functions as an RNAP antibacktracking factor by cleaving the nascent 3’ RNA that has extruded from the RNAP catalytic channel during backtracking(10, 55). Furthermore, GreA promotes the escape of RNAP from the promoter and suppression of promoter-proximal pausing during transcription initiation(56). To test whether GreA activity also contributed to RNAP release at the loci transcribing structured RNAs, we performed RpoB ChIP-seq of *B. subtilis* WT and a *ΔgreA* strain. We found that the *ΔgreA* strain only had 12 genes (and six TUs) with increased RpoB occupancy (Dataset S6, Fig. S7). Two of the six TUs transcribe regulatory RNAs and neither contained significant predicted secondary structure. These results suggest that unlike Mfd, GreA does not function in releasing RNAP from sites containing secondary structure. We also observed that in the *ΔgreA* strain, a total 469 genes exhibited less than two-fold RpoB occupancy (Dataset S6, Fig. S7). The high number of genes with decreased RpoB occupancy in the absence of GreA may be due to decreased efficiency of RNAP promoter escape during initiation and consequently decreased levels of elongating RNAP molecules, consistent with *in vitro* findings(56, 57).

### Mfd decreases expression at structured, regulatory RNAs

We tested the effect of Mfd on transcription at sites containing structured RNAs. We began by performing RNA-seq of WT and *Δmfd* strains in *B. subtilis*. Differential expression analysis between WT and *Δmfd* revealed a total of 378 genes with statistically significant lower RNA levels (Dataset S7). Consistent with our WT and *Δmfd* RpoB ChIP-seq results, we found that more genes were upregulated than downregulated in the *Δmfd* strain (240 genes upregulated compared to 138 genes downregulated) (Fig. S8 and Dataset S7). When comparing our RpoB ChIP-seq findings to RNA-seq, we found that of the 116 genes with greater than 2-fold RpoB ChIP-seq signal in the *Δmfd* strain, 30 of them directly showed increased expression, while none show decreased expression in the absence of Mfd. Analysis of genes with decreased expression in the *Δmfd* strain showed an equal decrease throughout the gene body, suggesting that Mfd suppresses full-length transcripts (Fig. S9). Because standard RNA sequencing protocols are often not suitable for accurate measurement of small RNAs(58), we wondered if there are additional genes with increased RpoB density in the *Δmfd* strain that have corresponding increases in transcription, but were not accurately detected in our RNA-seq analysis. We therefore directly measured RNA levels using qRT-PCR at three loci containing non-coding RNAs (the *trnY* locus, *txpa-ratA*, and *bsrH-asBsrH*), all of which show Mfd binding and increased RpoB signal in the *Δmfd* strain. We found that all three of these loci have increased gene expression in the *Δmfd* strain compared to WT (Fig. S10). To confirm that the *Δmfd* strain does not globally repress transcription, we performed qRT-PCR analysis on two control loci, *rpoB* and *yolA*, and found no difference in RNA levels between WT and the *Δmfd* strain (Fig. S10). These findings suggest that Mfd’s *in vivo* effects on RNAP lead to decreased transcription.

### Cells lacking Mfd are highly sensitized to toxin overexpression

Aside from decreased rates of evolution, there are few phenotypic defects that have been detected in the absence of Mfd, even upon exposure to DNA damage(17, 18, 24, 25, 59). We wondered whether the transcriptional regulation activity of Mfd at regions we detected were physiologically relevant. To address this, we focused on the highest structured regulatory RNAs which had altered RpoB density in the *Δmfd* strain and were also sites of Mfd association. These regulatory RNAs were present in two pairs of type I toxin-antitoxin (TA) loci in *B. subtilis:* the *txpA/ratA* locus and the *bsrH/as-bsrH* locus. Type I TA loci are characterized by the expression of a small toxic peptide and a noncoding RNA that neutralizes toxin expression by direct binding and either inhibiting translation or promoting degradation of the toxin mRNA(60). The cellular functions of type I TA loci remain unclear, but they have been proposed to be important for diverse aspects of physiology, including persister formation(61), biofilm formation(62), and prophage maintenance(63). Five type I TA loci have been identified in *B. subtilis*(60) - we found that three of these loci have both Mfd binding and a minimum of two-fold increase in RpoB density in the *Δmfd* strain, while a fourth locus, *yonT/as-YonT*, showed a significant increase in RpoB density in the *Δmfd* strain (Table S3).

Transcription regulation at type I TA loci is essential for cell survival as overexpression of type I toxins can be lethal(64). Based on our data, we hypothesized that cells lacking Mfd would be sensitive to toxin over-expression, similar to ethanol stress and heat shock(60). Because of the potential pleotropic effects, we chose to directly test our hypothesis by upregulating toxin expression in a controlled fashion. We therefore constructed an integrative plasmid containing the *txpA* toxin gene under an IPTG (isopropyl b-D-1-thiogalacto-pyranoside) inducible promoter in both WT and *Δmfd* strains, and performed cellular viability assays. We found that cells lacking Mfd are highly sensitized to both chronic (Fig. 6*A*) and acute (Fig. 6*C*) overexpression of TxpA. Cells lacking Mfd showed up to five orders of magnitude sensitivity to overexpression of this toxin. Similarly, we overexpressed the BsrH toxin in WT and *Δmfd* strains to test whether Mfd’s effect was conserved. Indeed, we see roughly four orders of magnitude sensitization of the *Δmfd* strain to overexpression of BsrH in both chronic (Fig. 6*B*) and acute (Fig. 6*D*) conditions. To test whether this effect was directly due to overexpression of BsrH or TxpA, we performed qRT-PCR analysis in our WT and *Δmfd* strains containing the overexpression constructs. We found that in both cases, toxin overexpression was increased by ~2-3-fold in the *Δmfd* strain (Fig. 6*E-F*). To confirm that the effects we observed were not specific to our integration site or promoter, we performed qRT-PCR analysis on strains containing an IPTG-inducible *lacZ* gene. We find no effect on *lacZ* transcript levels between WT and the *Δmfd* strain (Fig. S11). Interestingly, recent data has revealed that *Clostridium difficile* cells lacking Mfd have morphologic differences and viability defects that are thought to be related to toxin overexpression(65), underscoring the likely conserved and critical role that Mfd may play on the expression of such regulatory RNAs. Overall, our findings suggest that Mfd plays an important role in regulating transcription of TA loci in *B. subtilis*.

**Fig. 6.**
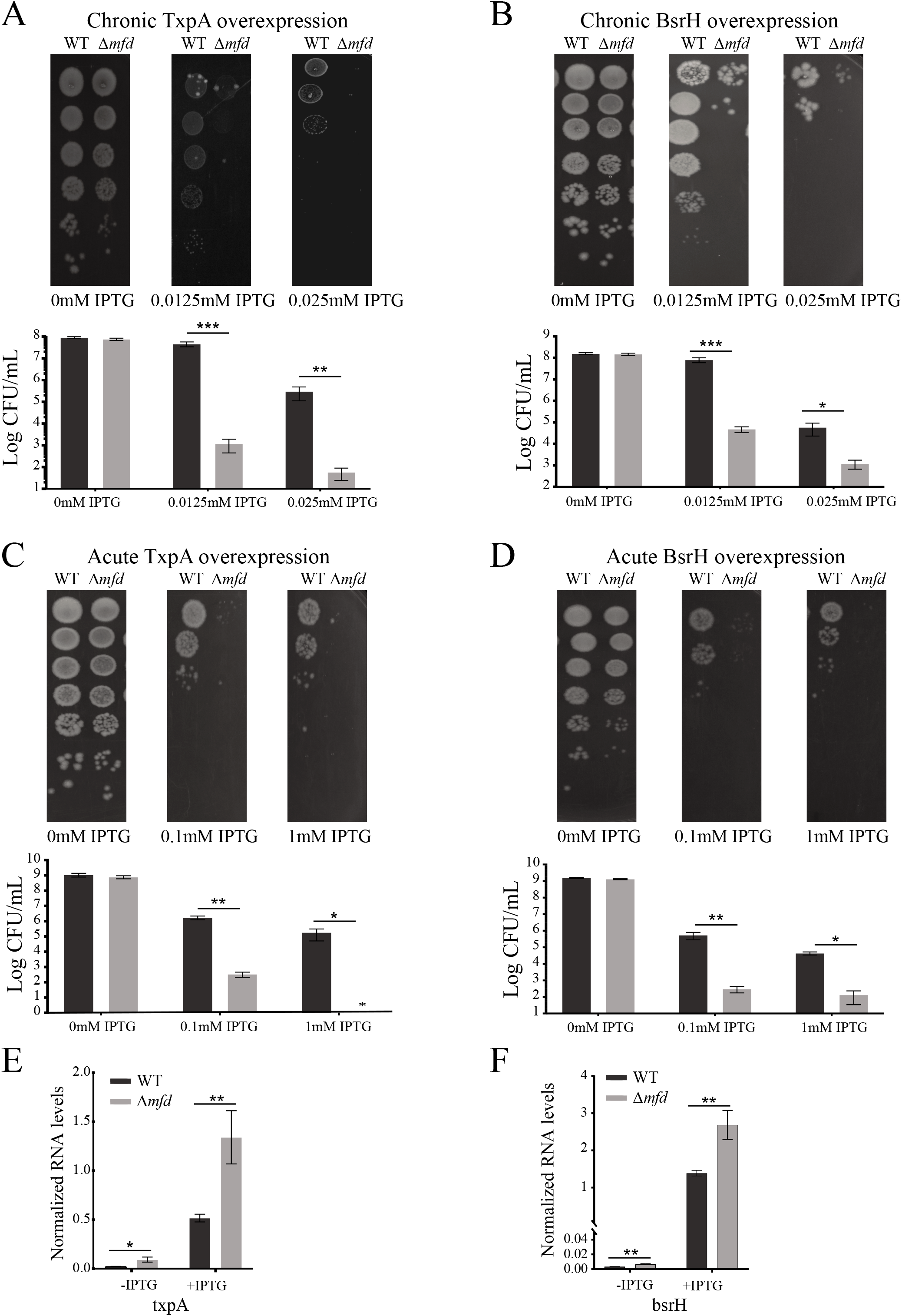
Transcriptional regulation by Mfd at toxin-antitoxin loci is critical for cell survival. Survival assays under chronic (A) and acute (C) overexpression of TxpA toxin (MFE Z-score = −6.77) in WT and *Δmfd* and survival assays under chronic (B) and acute (D) overexpression of BsrH toxin (MFE Z-score = −7.51) in WT and *Δmfd*. For all figures, representative images shown above, and quantification of data shown below. Error bars represent the SEM from at least three independent experiments. (E and F) qRT-PCR analysis of txpA and bsrH overexpression in WT and *Δmfd* strains. RNA values normalized to ribosomal RNA. Error bars represent the SEM from at least two independent experiments. Statistical significance was determined using a two-tailed Student’s T-test (*p<0.05, **p<0.01, ***p<0.001).

### Mfd promotes mutagenesis of the *txpA* gene

Mfd is an evolvability factor(17) and promotes mutagenesis under various conditions, promoting the development of antimicrobial resistance, and mutagenesis in general during nutrient starvation, stationary-phase growth, and replication-transcription conflicts(24, 25, 59, 66). Most of these studies utilize engineered reporter systems to study Mfd-mediated mutagenesis. It remains unclear what endogenous sites are prone to Mfd-mediated mutagenesis. Given our findings that Mfd is critical for transcriptional regulation at structured RNAs, we wondered if such sites were particularly prone to Mfd-mediated mutagenesis. We tested this hypothesis by performing Luria-Delbrück fluctuation assays(67) to select for toxin-resistant mutants and assessed the mutation rate at our ectopic *txpA* locus. We plated cells on a high concentration of IPTG (1mM) to ensure that only genetic revertants would produce viable colonies. We found that the mutation rate of *txpA* is nearly 7-fold lower in the *Δmfd* compared to the WT strain (Fig. S12). These differences are higher than previously reported mutation rate reductions in *Δmfd* strains(17, 59). We conclude that Mfd promotes mutagenesis at TA loci in *B. subtilis* and given our prior findings, it likely accelerates evolution at many of its endogenous targets.

## Discussion

In this work, we identified the endogenous targets of Mfd and how Mfd impacts RNAP association and transcription *in vivo*. We found that in both *E. coli* and *B. subtilis*, most of the sites where Mfd associates and modulates transcription contain highly structured regulatory RNAs. We found that RNAP modulation and transcriptional control by Mfd predominantly occur at regions of frequent RNAP pausing. Our experiments demonstrated that Mfd is important for maintaining cellular viability upon activation of TA systems that contain highly structured RNAs. We also showed that Mfd promotes mutagenesis at these regions. Based on these findings, we propose that Mfd is a RNAP co-factor which helps regulate transcription at many chromosomal regions, especially those with highly structured RNAs.

Though Mfd is non-essential under many conditions, it becomes critical for cell viability when TA systems are expressed. This suggests that there may be other situations when transcriptional regulation by Mfd is important. It is not unreasonable to expect that Mfd would be critical under conditions related to the functions of the genes where Mfd binds and regulates RNAP association. For example, Mfd targets a mannose utilization operon, suggesting that efficient growth on mannose may require transcriptional regulation by Mfd. By identifying the genomic regions where Mfd binds, our study has provided key information regarding the conditions under which Mfd function may become critical. Further work is needed to identify and characterize these conditions.

Though Mfd clearly reduces RNAP density at regions containing structured RNAs, it is not clear if this is due to RNAP rescue, early transcription termination, or a combination of both (Fig. 7). Though Mfd can rescue or terminate transcription, the decision about which activity will occur is thought to be related to the magnitude of the roadblock faced by RNAP, with more significant roadblocks favoring termination(10, 21, 22, 34, 68, 69). Additionally, recent work from Le et al. using *in vitro* translocase assays showed that Mfd rescues RNAP at pause sites, but that more severe obstacles to RNAP movement lead to eventual termination(23).

**Fig. 7.**
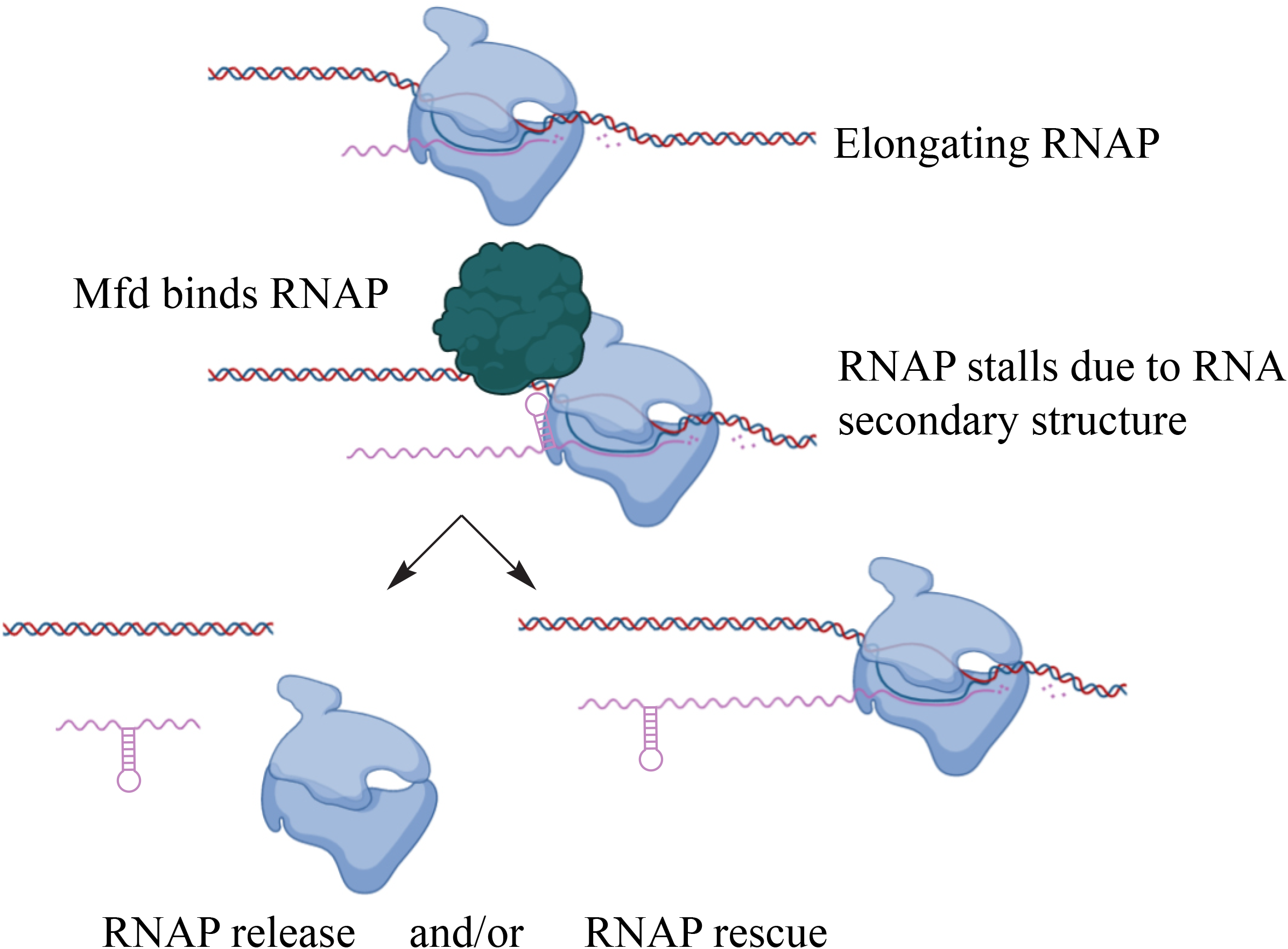
Model of Mfd activity at structured regulatory RNAs. During transcription, elongating RNAP (shown in blue) transcribes a highly structured RNA sequence. The formation of a structured RNA within the exit channel of RNAP can pause RNAP on DNA, leading to Mfd binding (shown in red). Two (non-mutually exclusive) models are shown to explain Mfd’s function once RNAP encounters a highly structured RNA. Bottom left: the arrested complex is recognized by Mfd, which releases RNAP from the template and in doing so represses transcription or bottom right: Mfd recognizes the arrested or stalled RNAP complex and functions as an elongation factor, accelerating RNAP release off of the DNA template.

Existing studies show that RNA secondary structure is capable of impeding RNAP processivity. For example, RNAP pausing can be promoted by the formation of stable RNA hairpin structures in the exit channel of RNAP and inhibit its movement(5, 70–72). Pausing via RNA secondary structure is a mechanism that can regulate gene expression at riboswitches, promote coupling of transcription and translation(73, 74), and is critical for the process of intrinsic transcription termination in bacteria(12, 75).

The severity of stalling at structured RNAs is likely dependent on multiple factors including the stability of the structure, length, and expression level. At certain sites, it is possible that such pausing may induce RNAP backtracking, but mechanistic studies suggest that it is more common for pausing to induce a “half translocated” state of RNAP(71) which inhibits its immediate processivity(76). To our knowledge, it has not been shown that RNA secondary structure is a significant enough roadblock to cause RNAP termination, suggesting that it is a generally less severe of an impediment to RNAP movement. These findings would favor a model whereby Mfd functions to promote RNAP rescue and elongation at structured RNAs, leading to the decreased RNAP levels as we observe in our ChIP studies. However, we cannot rule out the possibility that the regulatory RNAs identified in our studies promote a more severe impediment to RNAP and that decreased RNAP levels reflect RNAP termination by Mfd.

Given the wide range of biological functions regulated by structured RNAs, Mfd’s activity at the sites we identified is likely to have a broad range of phenotypic effects. For example, we identified multiple riboswitches containing TUs where Mfd binds and promotes RNAP release. These TUs are involved in many critical metabolic processes, ranging from beta-glucoside metabolism (*bglP-bglH-yxiE*)(77, 78), to the utilization of glycerol (*glpF-glpK* and *glpT-glpQ*)(79) to purine metabolism (*purEKBCSQLFMNHD*)(80, 81). We also identified Mfd binding and RNAP release at a locus containing a long cis-acting antisense RNA (*yabE*/S25) that is thought to play a role in cell wall maintenance(82). Lastly, amongst many other sites, we identified Mfd binding and RNAP termination at tRNA loci (the *trnY* locus in *B. subtilis* containing a highly structured RNA of unknown function). The physiological relevance of Mfd’s activity at these sites requires further investigation that is outside of the scope of this study.

Various mechanisms of transcription-associated mutagenesis (TAM) exist(83, 84). Based on our findings, we propose that the inherent structure of RNA may be an additional mechanism by which transcription promotes mutagenesis, at least partially through Mfd. Interestingly, RNA secondary structure has been reported to enhance mutation rates in replicating retroviruses(85), suggesting that evolution via RNA secondary structure may be a universal mechanism. In addition, a recent study by Thornlow et al.(86), using computational analyses, revealed that tRNAs have higher mutation rates relative to other parts of the genome(86). They also suggested that this phenomenon is linked to transcription. It is therefore quite possible that, at least in bacteria, the evolution of tRNA structures is mediated by Mfd. By promoting mutagenesis at sites of highly structured RNAs, Mfd may inherently alter the secondary structure encoded at the site of its activity, leading to novel or altered functions of the RNA. Additionally, non-coding RNAs are well-known to evolve very quickly(87), however, the mechanisms by which this occurs is unknown. Our results suggest that Mfd may contribute to the evolution of these regions via its mutagenic activity. Addressing this possibility would further contribute to our mechanistic understanding of how non-coding RNAs evolve.

## Materials and Methods

### Strain constructions

All strains and plasmids used and constructed in this study are listed in Table S4 and primers used are listed in Table S5. *B. subtilis* stains used in this study were derivates of the HM1 (JH642) parent strain(88). *E. coli* strains used were derivates of K-12 MG1655 (89).

Transformations into *B. subtilis* HM1 were performed under standard conditions as previously described(90). Plasmids used in this study were grown in *E. coli* DH5α. Plasmids were cloned using chemical transformations of competent *E. coli*. All plasmid purification was performed by growth of appropriate *E. coli* strain overnight at 37° C in Luria-Bertani (LB) medium supplement with the appropriate antibiotic and plasmids were subsequently purified using the GeneJet Plasmid Miniprep Kit (Thermo). Further details on strain construction can be found in the SI appendix.

### Growth conditions

For experiments in *B. subtilis* and *E. coli*, cultures were grown as described unless otherwise indicated. Cells were plated on LB supplemented with the appropriate antibiotic for isolation of single colonies. Overnight cultures from single colonies were grown at 37° C in LB at 260 RPMs and the following day cells were diluted back to OD600 0.05 and grown to exponential phase (OD600 0.3-0.5) before harvesting. For acute rifampicin ChIP experiments, cultures were grown in identical fashion until they reach OD600 0.3-0.5 and rifampicin was added at a concentration of 100μg/mL for 5 minutes before harvesting.

### ChIP-seq and ChIP-qPCR experiments

For *B. subtilis* Mfd ChIP experiments, c-Myc mouse monoclonal antibody (clone 9E10) was used (Thermo). For *E. coli* ChIP experiments, custom polyclonal *E. coli* Mfd rabbit antisera was produced and purified by Covance. ELISA titers were used to confirm Mfd titers from antisera. For RpoB experiments, RNA polymerase beta mouse monoclonal antibody (clone 8RB13) was used (Thermo).

ChIP experiments were performed as previously described(91, 92). Briefly, Cells were grown to exponential phase as previously described and crosslinked with 1% formaldehyde v/v. After 20 minutes at room temperature, 0.5M final concentration of glycine was added and cells were pelleted, washed in cold 1x PBS and pelleted again. Cells were resuspended in solution A (10 mM Tris pH 8.0, 10 mM EDTA, 50 mM NaCl, 20% sucrose), supplemented with 1 mg/ml lysozyme and 1 mM AEBSF at 37° C for 30 minutes. 2x IP buffer (100 mM Tris pH 7.0, 10 mM EDTA, 300 mM NaCl, 20% triton x-100), supplemented with 1mM AEBSF, was then added and lysates were incubated on ice for 30 minutes. Cell lysates were sonicated four times at 30% amplitude for ten seconds using a Fisher sonic dismembrator (Fisher FB120). Lysates were centrifuged at 8000 RPMs for 15 minutes at 4° C. The supernatant was transferred into new microfuge tubes.

ChIP lysates were split into a total DNA input control (40μl of lysate) and immunoprecipitation (IP) (1mL of lysate). For Mfd ChIP experiments in *B. subtilis* 12μl anti-c-Myc antibody was added to the IP samples. For Mfd ChIP-seq experiments in *E. coli*, 4 μl of native *E. coli* anti-Mfd antibody was used. 2μl of anti-RpoB antibody was added for RpoB ChIPs in both *B. subtilis* and *E. coli*. IP lysates were rotated overnight at 4° C. The following day, 30μl Protein A sepharose beads (GE) were added to the IP samples and rotated for one hour at room temperature. Beads were then pelleted with centrifugation at 2000 RPMs for one minute. Supernatant was decanted and beads were subsequently washed six times with 1x IP buffer and one time with 1x TE pH 8.0. Beads were then pelleted and resuspended in 100μl of elution buffer (50mM Tris pH 8.0, 10mM EDTA), and 1% SDS and incubated at 65° C for 10 minutes. Beads were pelleted by centrifugation and supernatant was transferred to a new microfuge tube. A second round of elution was performed by resuspension of beads in 150μl of elution buffer II (10mM Tris pH 8.0, 1 mM EDTA, 0.67% SDS). Beads were pelleted and supernatant was transferred to microfuge tube containing eluate from the first elution. IP samples were then incubated overnight at 65° C. The following day, proteinase K was added at a final concentration 0.4 mg/mL and samples were incubated for two hours at 37° C. Purification was performed by using the GeneJet PCR Purification Kit (Thermo).

Library preparation for ChIP-seq was performed using the Nextera XT DNA Library Prep Kit (Illumina) according to manufacturer’s instructions. For ChIP-quantitative PCR (qPCR), Sso Advanced Universal SYBR Green Supermix (BioRad) was used according to manufacturer’s instructions.

### NET-seq analysis

Data from Larson et al. (41) was analyzed for determination of average NET-seq read counts in addition to identification of pause sites. Data was collected from Gene Expression Omnibus (accession number GSE56720) and plotted as a wiggle (.wig) file on the *B. subtilis* 168 genome. Details of pause site determination are defined in the supplementary methods of Larson et al. (41). In order to calculate the average read count for each gene, gene coordinates were determined and NET-seq signal was averaged across these nucleotide positions. Average NET-seq signal for each gene was calculated from the strand (plus or minus strand) that had the highest NET-seq signal. The number of pause sites identified for each gene of interest was normalized to gene length.

### PARS-seq analysis

PARS data was kindly provided by Zoya Ignatova. Original PARS score data was refined to remove duplicate data points. All positive PARS scores were then plotted as a wiggle (.wig) file on the *E. coli* K-12 MG1655 genome sequence (Genbank: U00096.2). The average PARS score for each annotated feature in the *E. coli* chromosome was calculated using custom scripts.

### Whole-genome sequencing analysis

ChIP-seq and RNA-seq samples were sequenced using the Illumina Nextseq 500/550 Sequencing system at the University of Washington Northwest Genomics Center and the VANTAGE Sequencing Core at Vanderbilt University. After sequencing, sample reads from *B. subtilis* were mapped to 168 genome (accession number: NC_000964.3) and from *E. coli* to the MG1655 genome (accession number: NC_ U00096.2) using Bowtie2(93). For data visualization, SAMtools was used to process SAM files(94) to produce wiggle plots(95). Wiggle files from all ChIP samples were normalized to input samples (total input DNA subtracted from the ChIP signal). For quantification of ChIP-seq and RNA-seq samples, BAM files were processed by the featureCounts program to determine read counts per gene(96). To determine differential RNA-seq expression (see supplementary materials and methods for RNA-seq experiments) and differential ChIP-seq binding, read counts were analyzed by DEseq2 software(97). In order to determine correlations between RpoB ChIP binding and Mfd ChIP, read counts generated by featureCounts were divided by the total number of sequencing reads per sample. ChIP samples were then divided by input samples and log2 normalized.

### Quantitative RT-PCR assays

For quantification of RNA using qRT-PCR, *B. subtilis* cultures were grown to exponential phase as described in the growth conditions sections. For experiments containing IPTG-inducible constructs (P_spank_-txpA, P_spank_-bsrH, and P_spank_-lacZ), expression was induced with 1mM IPTG for 5 minutes and RNA was subsequently extracted as described in the RNA-seq experiments section (see supplementary materials and methods). Subsequently, 1μg of total RNA was treated with DNaseI (Thermo) for one hour at 37° C. DNase denaturation was performed with addition of 10mM EDTA and incubation at 65° C for 10 minutes. cDNA generation was performed using the iScript Supermix (BioRad), according to the manufacturer’s instructions. Quantitative PCR (qPCR) was performed using the Sso Advanced Universal SYBR Green Supermix (BioRad), according to manufacturer’s instructions. For normalization of qRT-PCR, primers to *B. subtilis* rRNA was used.

### Cell survival assays

For chronic survival assays, strains were struck out on LB agar plates and *B. subtilis* cultures were grown in 2mL LB until they reached an OD600 of 0.5-1.0. All cultures were normalized to OD600 0.3 and serial dilutions were performed in 1x Spizizen’s salts. 5μl of cells were plated on control plates containing LB agar only and LB agar plates containing the designated concentration of IPTG (see figure legends for concentrations). Plates were grown at 30° C overnight and CFUs were enumerated the following day.

For acute survival assays, cultures were grown in 2ml until they reached an OD600 of 0.5-1.0 and then diluted back to OD600 0.05. Either 1mM or 0.1 mM IPTG was then added and cells were grown for approximately 60 minutes (OD600~0.3). Cells were subsequently washed two times with 1x Spizizen’s salts to remove residual IPTG and were serially diluted. 5μl of cells were plated on LB agar and plates were grown at 30° C overnight for CFU enumeration. For both chronic and acute survival assays, images were taken using the BioRad Gel Doc XR+ Molecular Imager.

## Supporting information

AllSup

Datatset S4

## Acknowledgements

We thank Patrick Nugent and Ankunda Kariisa for assistance with strain construction and bacterial 2-hybrid assays, Kevin Lang for helpful discussions, Sirena Tran for assistance with *E. coli* ChIP-seq experiments, Zoya Ignatova for PARS data set, Charlie Lee at the University of Washington and the VANTAGE genomics core at Vanderbilt for assistance with sequencing experiments. This work was supported by the NIH National Institute of Allergy and Infectious Diseases AI127422 and Training Program in Environmental Toxicology T32 ES007028 to K.B.

## Footnotes

Author contributions: M.N.R., C.M., and H.M. conceived of and designed experiments. M.N.R., C.M., K.B. performed experiments. M.N.R., C.M., and H.M analyzed data and wrote the paper.

To whom the correspondence should be addressed.

The authors declare no conflict of interest

