## Supplementary material for "Mfd regulates RNA polymerase association with hard-to-transcribe regions *in vivo*, especially those with structured RNAs": AllSup

**This PDF file includes:**

Expanded Materials and Methods
Figures S1 to S12
Tables S1 to S5
Legends for Datasets S1 to S7
SI References

### Expanded Materials and Methods

#### Detailed strain construction

In order to construct marker-less point mutation (L522A) of Mfd in *B. subtilis*, the pminiMAD2 plasmid was used as previously described (1). Briefly, HM2916 was constructed by transforming pHM707 into HM1 and grown at in LB broth containing MLS antibiotics at 22°C, the permissive temperature. *B. subtilis* strains were then incubated for 12 hours at 42°C while maintaining MLS selection. Cells were serially diluted and passaged multiple times at 22°C. Individual colonies were plated on LB plates with or without MLS to identify colonies which were MLS sensitive and had evicted the plasmid.

HM2769 was constructed by transforming pHM430 and pHM439 into HM2747. HM2771 was constructed by transforming pHM430 and pBRα into HM2747. HM2773 was constructed by transforming pHM439 and pACλCI into HM2747. HM2965 was constructed by transforming pHM431 and pHM439 into HM2747. HM2932 was constructed by transforming the HM2916 plasmid into HM1.

HM3157 was constructed using the transformation of SOE PCR product into HM1. First, Mfd-myc amplicon was generated using primers HM3759 and HM3760 and HM1 genomic DNA as a template in order to add a 1x myc sequence to the Mfd gene. Erm resistance cassette was amplified using pCAL215 plasmid DNA as a template and primers HM3854 and HM3969. These two respective amplicons were used as templated to generate a PCR SOE product using primers HM3759 and HM3854.

HM3808 was constructed by transformation of HM712 genomic DNA into HM1. HM3933 was constructed by transformation of HM1333 genomic DNA into HM1451. HM3947 was constructed by transforming HM3157 genomic DNA into HM2916. HM3986, HM3988, and HM3990 were constructed by transforming plasmid pHM676 into *E. coli* DH5α, HM1, and HM2521, respectively. HM4002, HM4003, and HM4004 were constructed by transforming plasmid pHM682 into *E. coli* DH5α, HM1, and HM2521, respectively.

#### Detailed plasmid construction

pHM430 was built using Gibson cloning from pACλCI-β-flap backbone (BamHI/NotI digested) and *B. subtilis* Mfd amplicon (AA492-AA625) amplicon with stop codon added (using primers HM3286 and HM3287). pHM431 was built using site-directed mutagenesis of pHM430 using primers HM3540 and H3541. pHM439 was built using Gibson cloning from pBRα-β-flap backbone (BamHI/NotI digested and *B. subtilis* rpoB amplicon (AA21-AA131) with stop codon added (using primers HM3292 and HM3293).

pHM676 was built using digestion of pDR111 with sphI and subsequent ligation of a PCR amplicon generated with primers HM5418 and HM5419 and HM1 genomic DNA as a template.

pHM682 was built using digestion of pDR111 with sphI and subsequent ligation of a PCR amplicon generated with primers HM5462 and HM5463 and HM1 as a template.

pHM707 was built using digestion of pminiMAD2 with kpnI and bamHI and subsequent ligation of a PCR amplicon with primers HM1004 and HM1005 with HM1 genomic DNA as a template. Mutations were subsequently introduced via site-directed mutagenesis using primers HM3540 and HM3541.

**Western blot assay**

Exponentially growing cultures were centrifuged, resuspended in Tris/Salt buffer (50 mM Tris-HCl pH 8, 300mM NaCl), and pelleted. Cell lysis buffer (10mM Tris-HCl pH7, 10mM EDTA, .1mM AEBSF, .1mg/ml lysozyme) was added and samples were incubated at 37° C for 15 minutes. SDS loading buffer was added to samples and 20µl was loaded onto Mini-PROTEAN TGX Precast Gels (BioRad) and run in Tris/SDS/Glycine running buffer in a Mini-PROTEAN Electrophoresis Cell (BioRad) at 200V for 40 minutes. Transfer was performed using the Trans-Blot Turbo Transfer System (BioRad). Anti-c-Myc antibody (1:5000 dilution) was added and blots were incubated overnight at 4° C. Anti-mouse antibodies (Li-Cor) (1:15000 dilution) was added and blot was imaged using the Odyssey CLx imaging system (Li-Cor).

**Bacterial 2-hybrid assays**

Bacterial 2-hybrid assays were performed as previously described (2). Briefly, RNAP interacting domains of *B. subtilis* Mfd (WT and L522A) and the Mfd interacting domain of RpoB were fused to Lambda repressor the N-terminal domain of *E. coli* RNAP alpha subunit. Fusions were subsequently transformed into a strain of *E. coli* containing the lambda operator sequence upstream of a luciferase reporter gene. In order to measure relative light units (RLUs), *E. coli* strains were grown overnight at in LB + 20mM IPTG at 30° C. The following day, cells were diluted 1:100 into LB+20mM IPTG and growing until OD600 ~2.0. Measurement of RLUs was performed using the Nano-glo substrate (Promega), according to the manufacturer's instructions. Luminescence was measured using the SpectraMax M3 96-well plate reader.

**RNA-seq experiments**

*B. subtilis* cultures were grown to exponential phase as previously described and harvested by addition of 1:1 volume 100% cold methanol and centrifugation at 5000 RPMs for five minutes. Samples from WT and *Δmfd* were normalized by adding an equal volume of cells across all samples. Cell pellets were subsequently lysed in TE and lysozyme (20mg/mL) and purified using the GeneJet RNA Purification Kit (Thermo). A total of 1ug of RNA from each sample was used to performed library preparation for RNA-seq, performed using the Scriptseq Complete Kit (Bacteria) from Illumina, according to manufacturer's instructions. To determine differential RNA-seq expression between WT and *Δmfd*, read counts were analyzed by DEseq2 software (3). Details of inclusion criteria to define transcription differences are described in Dataset 7 legend below. RKPM ($log_2$ normalized) plots were generated for visualization purposes by normalizing the total number of mappable reads at each gene to total number of sequencing reads and to the gene length. RKPM values were then log normalized and averaged across two independent replicate experiments for both WT and *Δmfd*.

**Mutation rate analysis**

Luria-Delbrück mutation rate assays were performed as previously described (4). *B. subtilis* were grown on LB plates overnight and cultures from single colonies and subsequently grown in LB media at 37° C at 260 RPMs to exponential phase growth (OD600= .5). Cells were diluted back to OD600= 0.0005 and dispended into 2mL parallel cultures containing 2mL LB and grown in the same conditions until OD600=0.5. To identify mutants that were resistant to toxin overexpression, 100µl of each 2ml culture was plated on LB plates containing 1mM IPTG.  Cells were serially diluted and plated on LB for CFU enumeration. Colonies were quantified after growth overnight at 37° C for IPTG plates and 30° C for LB plates. Mutation rates were calculated using the Ma-Sandri-Sarkar Maximum Likelihood method (5).

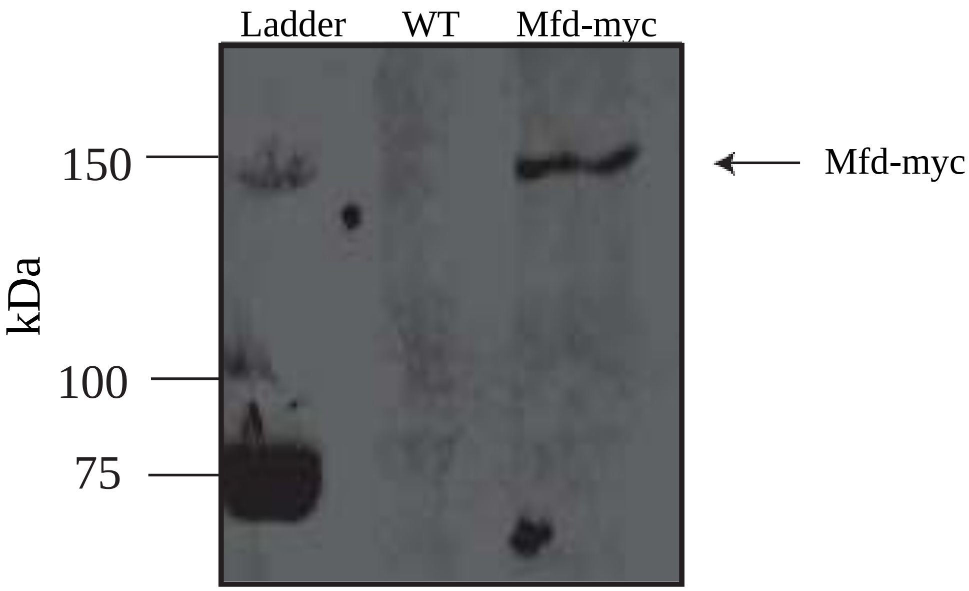


**Fig. S1. Western blots of *B. subtilis* Mfd-myc**

Western blot of *B. subtilis* WT and Mfd-myc. Anti-c-Myc antibody and anti-GFP antibody was used to probe blot.

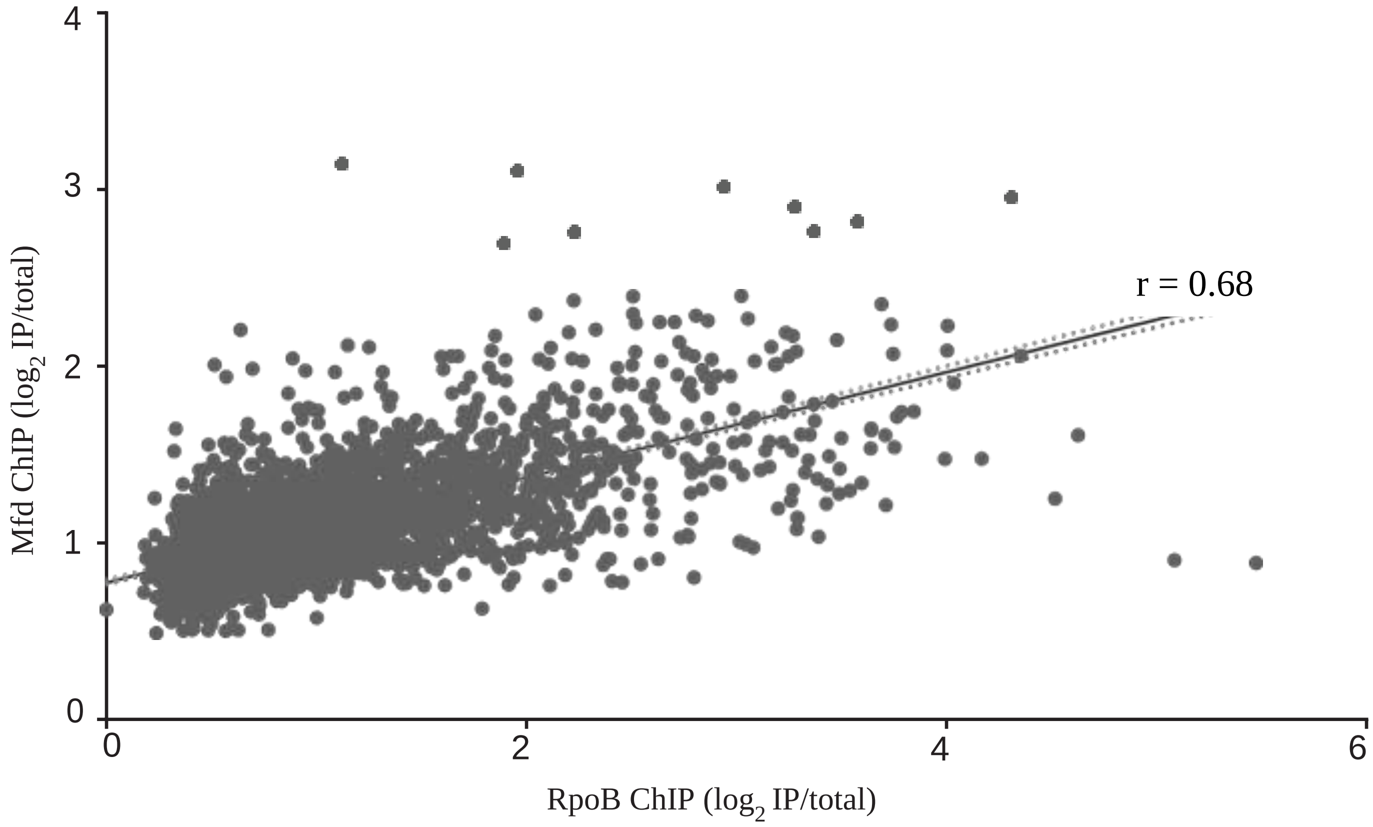


**Fig. S2. *B. subtilis* Mfd and RpoB ChIP-seq are correlated**

Linear regression analysis comparing binding of Mfd and RpoB at each gene in *B. subtilis*. Mfd-myc ChIP-seq (from Mfd-myc tagged *B. subtilis*) and RpoB ChIP-seq (from WT *B. subtilis*) read counts were determined for each gene in *B. subtilis* and normalized as described in Figure 2. Pearson's correlation coefficient for *B. subtilis* Mfd and RpoB = 0.68. Dotted lines represent 95% confidence interval.

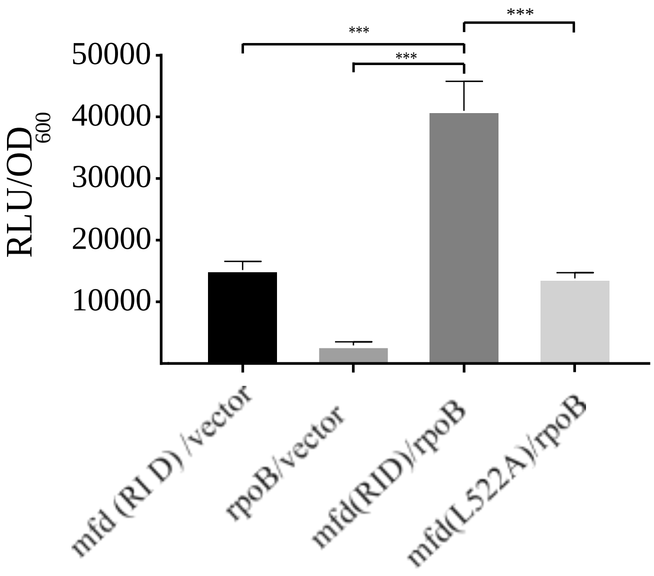


**Fig. S3. Bacterial two-hybrid assay exhibits abrogated binding between *B. subtilis* MfdL522A and RpoB.**

Disruption of Mfd L522 in *B. subtilis* abrogates interaction with RpoB. The interacting domains of RpoB and Mfd were cloned into a luciferase based bacterial 2-hybrid assay. Interactions between RpoB and Mfd and an MfdL522A mutant were measured, along with appropriate empty vector controls. Interactions were measured using luminescence and normalized to OD600. Data is from at least two independent experiments and error bars indicate standard deviation. Two-tailed students T-test was used to determine statistical significance (***p-value <0.001).

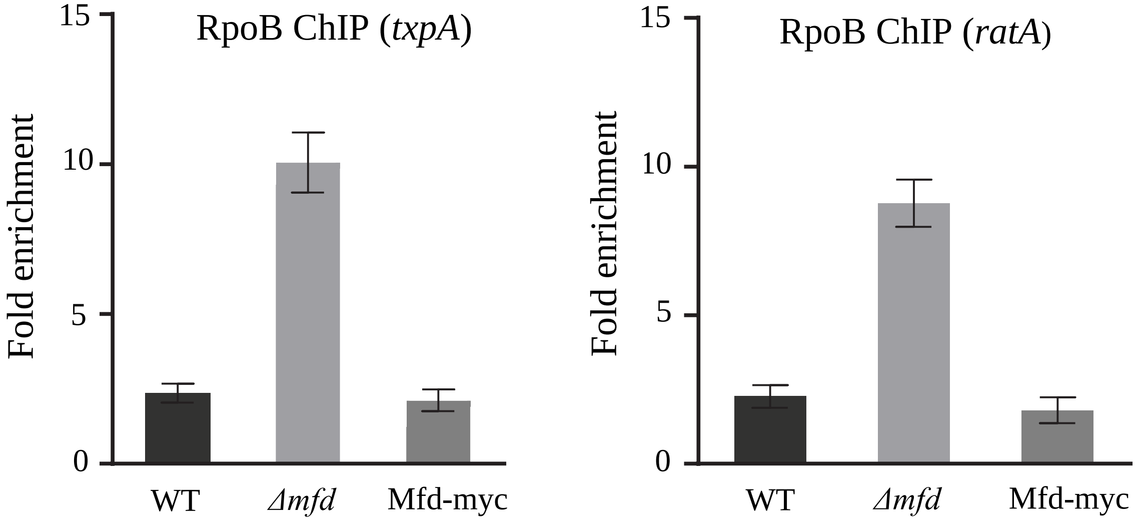


**Fig. S4. RpoB ChIP-qPCR corroborates ChIP-seq results and confirms functionality of Mfd-myc tagged strain**

Bar graph showing the normalized ChIP-qPCR levels for two genes (txpA- left and rataA- right), which shows increased RpoB signal in *Δmfd* via ChIP-seq analysis. Data collected from three different *B. subtilis* strains: (WT- black, *Δmfd* – light grey, Mfd-myc – dark grey). RpoB levels normalized to control locus *yhaX*. Data is from at least two independent experiments and error bars represent standard error of the mean (SEM).

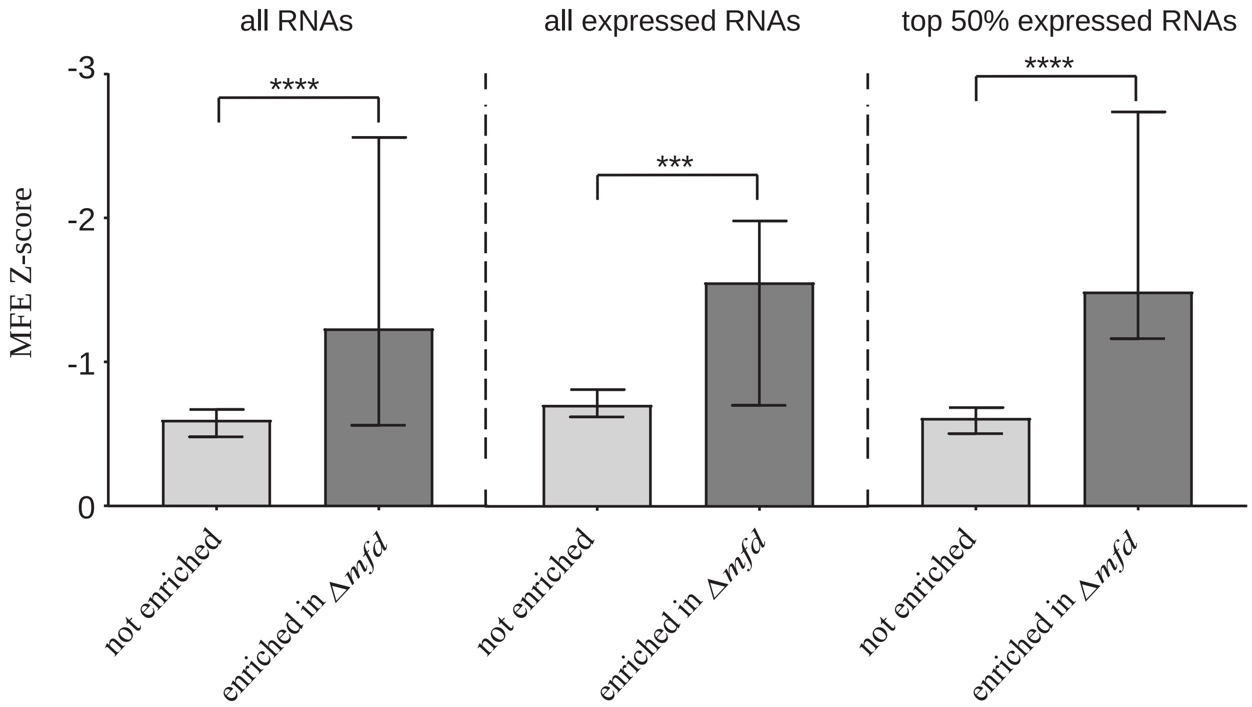


**Fig. S5. Transcription units with Mfd binding and increased RNAP density in *Δmfd* are enriched for structured regulatory RNAs**

Bar graph showing the median minimum free energy (MFE) for regulatory RNAs in *B. subtilis*. Grey bars represent regulatory RNAs within TUs that have no observed change in RpoB density between WT and *Δmfd* and black bars represent TUs that have increased RpoB density in *Δmfd* and are also bound by Mfd. Data is stratified by transcription levels, with all regulatory RNAs (expressed and non-expressed) shown in the left bars graphs, all expressed RNAs shown in the middle bar graphs, and the top 50% of expressed RNAs in the right. represent 95% confidence intervals. Statistical significance was determined using the nonparametric Mann-Whitney test for two population medians (***p<0.001, ****p<0.0001)

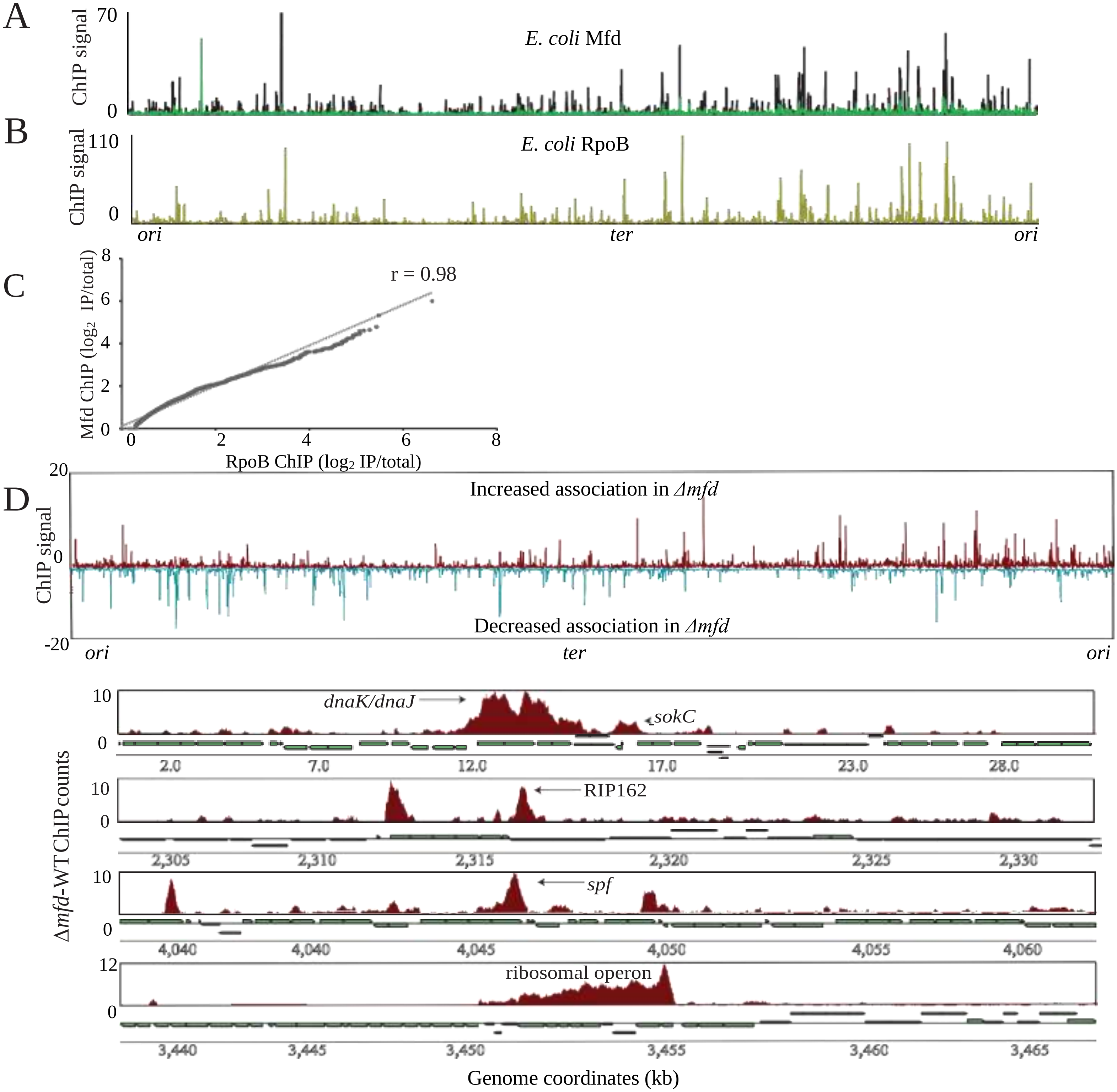


**Fig. S6. Mfd's effect on RNAP at sites of structured regulatory RNAs is conserved in *E. coli***

(A) ChIP-seq plot of Mfd in WT (dark blue) and *Δmfd* (green) *E. coli* using native antibody. (B) ChIP-seq plot of WT *E. coli* RpoB. Plots averaged from at least two independent experiments (C) Linear regression comparing binding of Mfd and RpoB at each gene in *E. coli*. Read counts were determined for each gene and log normalized as described in Figure 2. Dotted lines represent 95% confidence interval. (D) RpoB ChIP-seq plots showing regions of RpoB enrichment in *Δmfd*. Top half of graph (red) reflects normalized RpoB ChIP-seq read counts where *E. coli Δmfd* had increased signal relative to WT. Bottom half of graph (blue) reflects RpoB ChIP-seq read counts where *Δmfd* had decreased signal relative to WT *E. coli*. Four zoomed in plots (30kb window) below show ChIP signal at four sites thought to contain regulatory or structured RNAs (*sokC*, *spf*, RIP162, ribosomal operons).

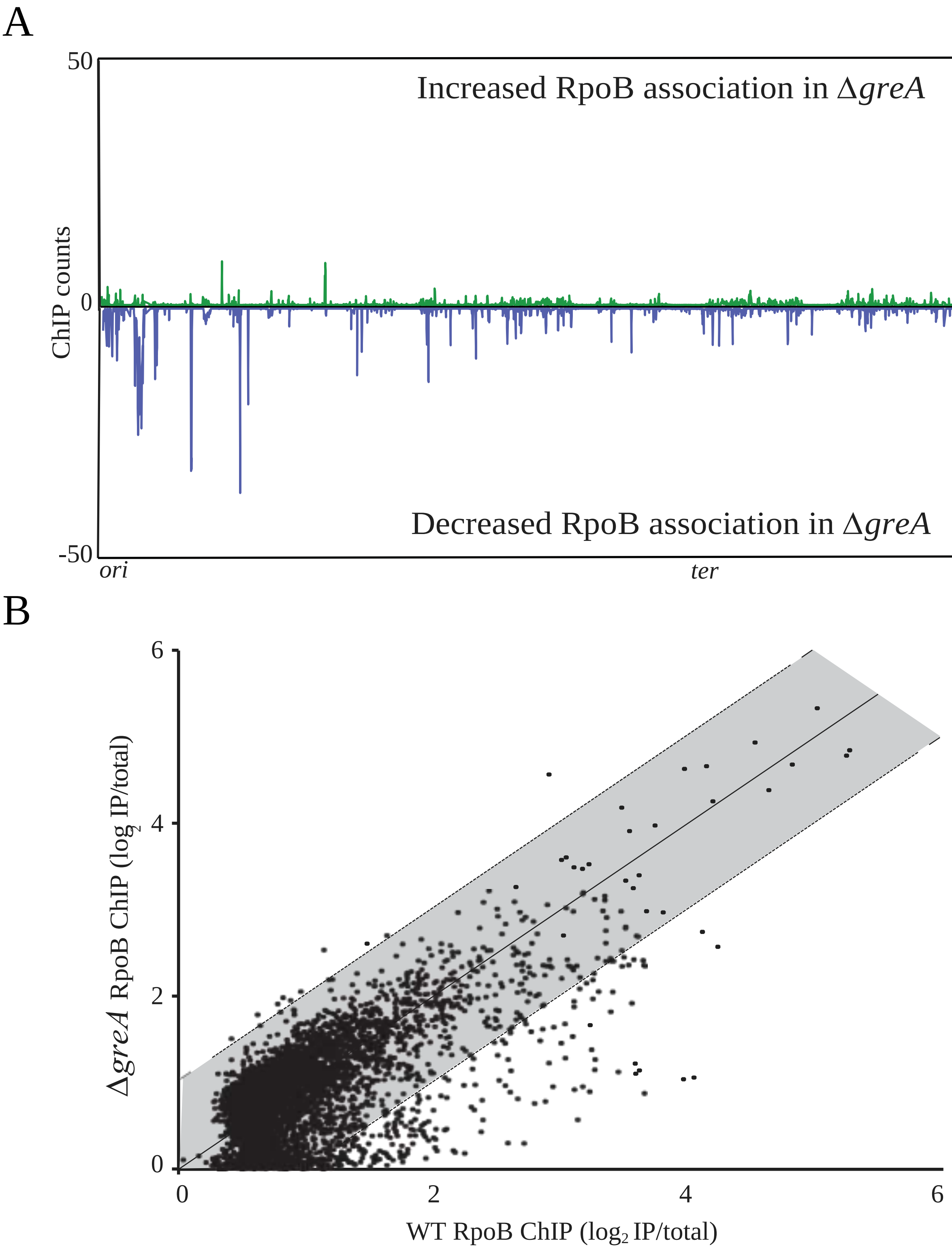


**Fig. S7. GreA does not alter transcription at sites containing regulatory RNAs**

RpoB ChIP-seq plots showing regions of RpoB enrichment in *ΔgreA* relative to WT. (A) Top half of graph (green) reflects normalized RpoB ChIP-seq counts where *B. subtilis ΔgreA* increased signal relative to WT. Bottom half of graph (blue) reflects RpoB ChIP-seq read counts where *ΔgreA* had decreased signal relative to WT. Plots averaged from at least two independent experiments. (B) Scatter plot of WT and *ΔgreA* RpoB ChIP-seq. Quantification of ChIP signal was performed as described in Figure 2B.

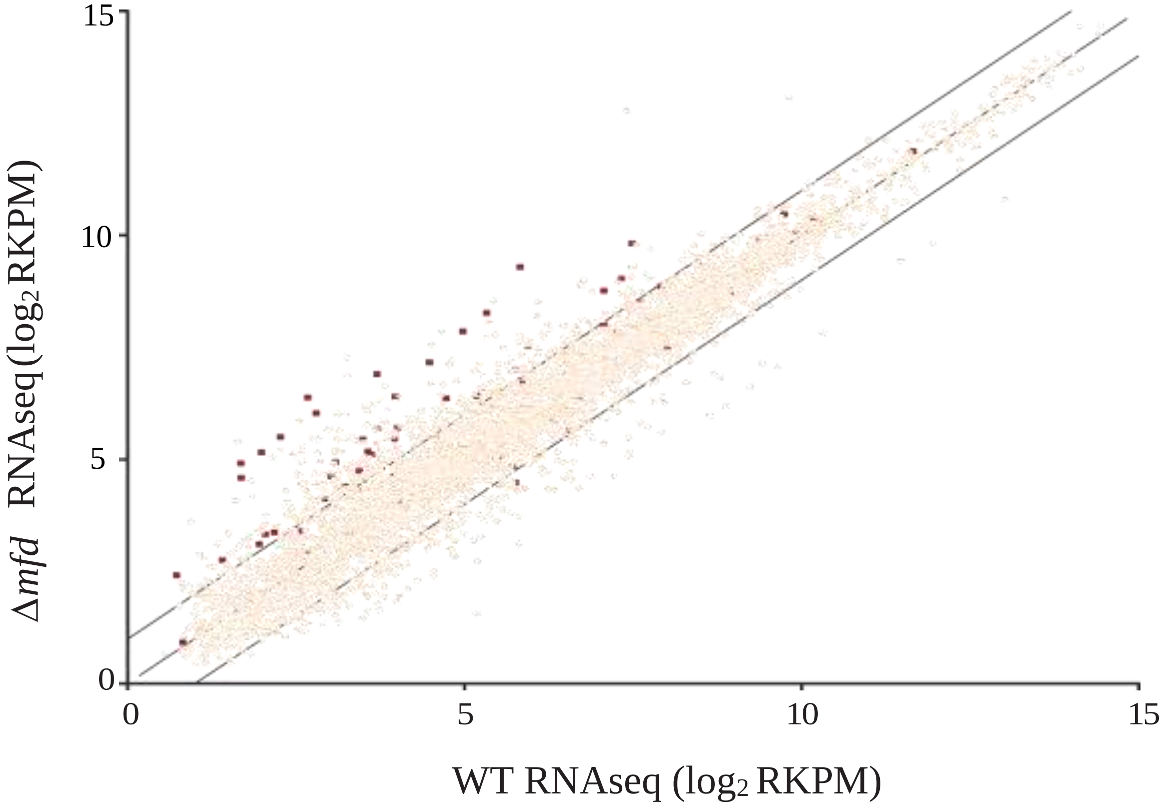


**Fig. S8. Mfd promotes decreased transcription at sites of structured regulatory RNAs**

Scatter plot of RNA-seq. Data points represent the expression level of each gene in *B. subtilis* in WT and *Δmfd* strains. Scatter plot represents expression level calculated using $\log_2$ normalized read per kilobase per million reads (RPKM)(13), from at least two independent experiments. Genes with increased RpoB occupancy in *Δmfd* shown as red squares.

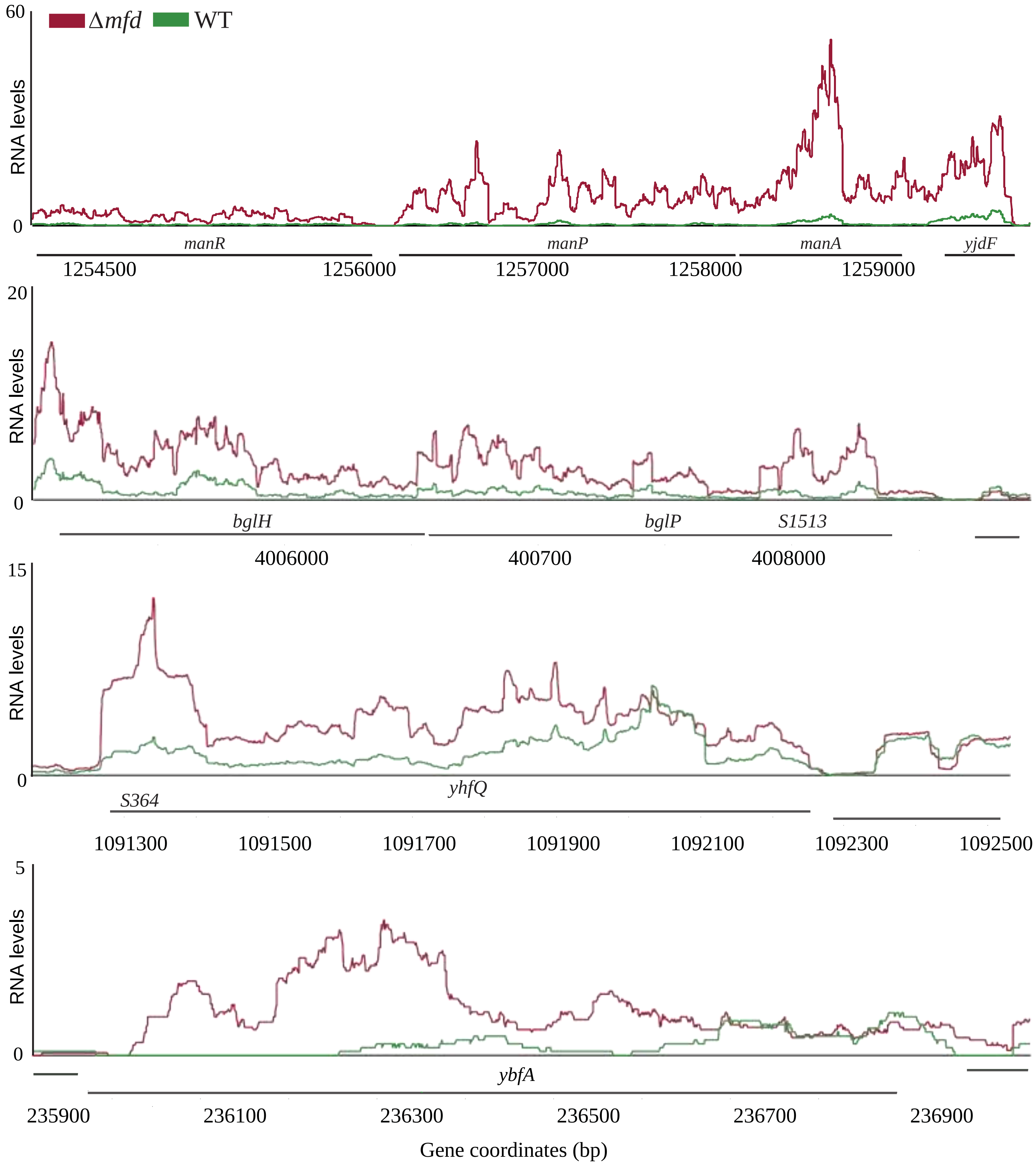


**Fig. S9. Expression of full-length transcripts repressed by Mfd**

RNA-seq plots showing transcription level at four representative loci with increased expressed in *Δmfd* (red) relative to WT (green).

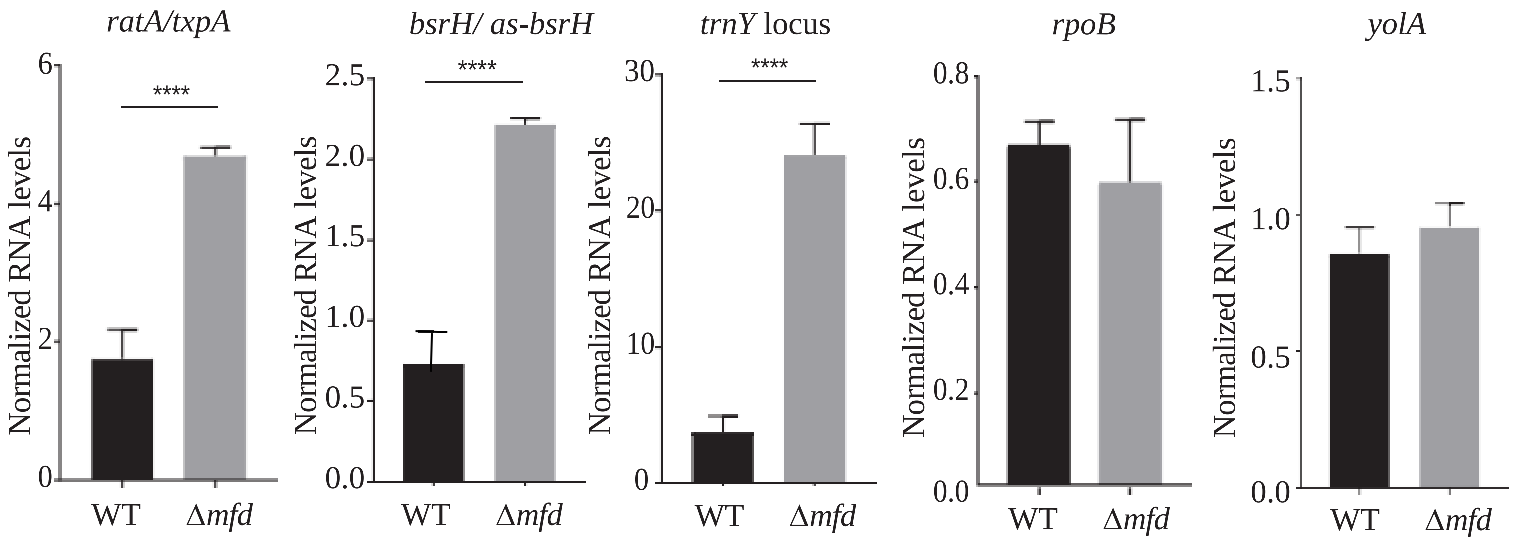


**Fig. S10. Mfd decreases transcription levels at select structural regulatory RNAs**

qRT-PCR analysis of three regions with increased RNAP occupancy in *Δmfd*: *txpA/ratA*, *bsrH/as-bsrH*, and the *trnY* locus (right), in addition to two control loci (*rpoB* and *yolA*). RNA values normalized to ribosomal RNA. Error bars represent the SEM from at least two different experiments. Statistical significance was determined using a two-tailed Student's T-test (****p<0.0001).

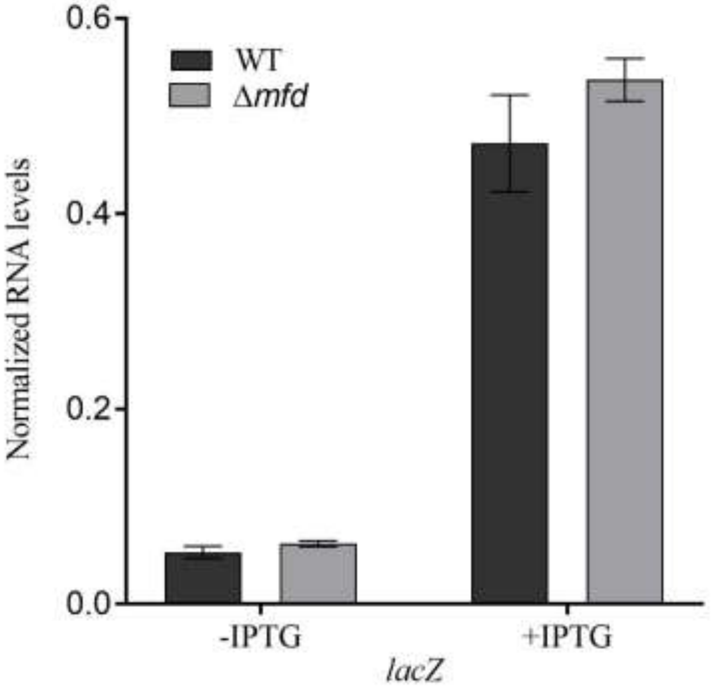


**Fig. S11. Mfd does not alter transcription of control reporter gene**

qRT-PCR analysis of *lacZ* overexpression under IPTG control, in WT and *Δmfd* strains. RNA values normalized to ribosomal RNA. Error bars represent the SEM from at least two independent experiments.

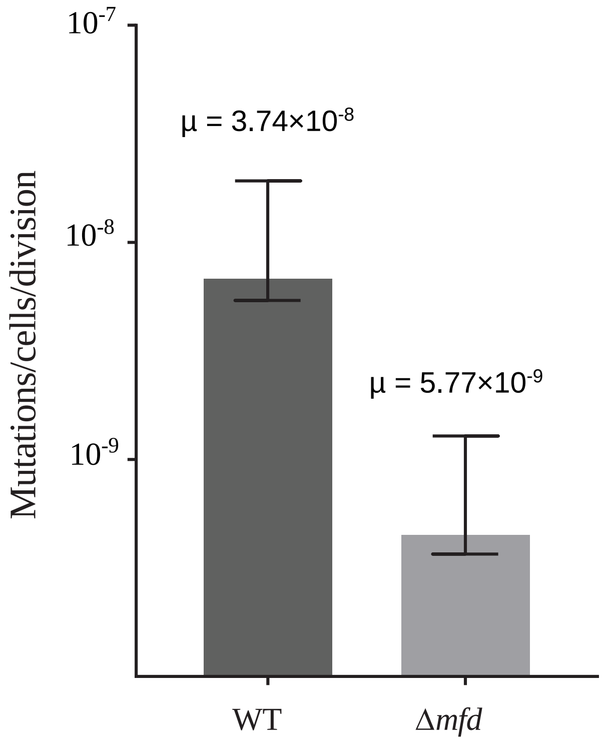


**Fig. S12. Mfd promotes mutagenesis at toxin-antitoxin loci**

Mutation rate analysis of WT (black) and of *Δmfd* (grey) strains containing ectopic TxpA overexpression ($P_{spank(hy)}$-*txpA*). At least 24 replicates were performed for each strain, from at least three independent experiments. Error bars are 95% confidence intervals.

| TUs with Mfd binding and increased RpoB in *Δmfd* | regulatory RNA category |
|---|---|
| *manP-manA-S439-yjdF* | riboswitch, intergenic |
| *S442-yjdH-S441-yjdG* | 5’ UTR, intergenic |
| *S81-ybeF-ybfA-ybfB* | 5’ UTR |
| *txpA (as ratA)* | asRNA, ncRNA |
| *bsrH (as bsrH)* | asRNA, ncRNA |
| *S782-S783-yopT* | 5’ UTR |
| *S345-S346-yhaX* | independent transcript |
| *S823-ilvD* | 5’ UTR |
| *yabE (S25 asRNA)* | asRNA, ncRNA |
| *yrrT-mtnN-S1033-mccA-mccB-yrhC* | Intergenic |
| *S1434-maeE* | 5’ UTR |
| *S1552-S1553-walR-walK-walH-walI-walJ-htrC* | 5’ UTR |
| *S655-S654* | independent transcript |
| *yhfO-yhfQ-S364-yhfP* | intergenic |
| *S460-mhqA* | 5’ UTR |
| *bsrG (SR4 asRNA)* | asRNA, ncRNA |
| *S438-yjdB-S437* | 5’ UTR, 3’ UTR |
| *S27-rnmV-ksgA* | 5’ UTR |
| *S1123-nifZ-thiI-sspA* | 5’ UTR |
| *S1487-S1486-cydA-cydB-cydC-cydD* | 5’ UTR |
| *S492-clpE* | 5’ UTR |
| *manR* | none |
| *ndoAI-ndoA asRNA(S163-S164-S165)* | asRNA |
| *S811* | independent transcript |
| *trnY* locus | tRNA, intergenic |
| *alaR-alaT-S1201* | 3’ UTR |
| *S1175-S1174-mntA-mntB-mntC-mntD* | 5’ UTR |
| *S1427-S1426-atpI-atpB-atpE-atpF-atpH-atpA-atpG-atpD-atpC* | 5’ UTR |
| *S895-yqxK* | 5’ UTR |
| *S1513-bglP-bglH-yxiE* | 5’ UTR, riboswitch |
| *S321-glpF-glpK* | 5’ UTR, riboswitch |
| *mhqN-mhqO-mhqP* | none |
| *S1203-yugF* | 5’ UTR |
| *spoVM* | none |
| *S966-sdA (asRNA s965)* | 5’ UTR, asRNA |
| *polY2-yqjX-S898-S897-yqjY-yqjZ-yqkA-yqkB-yqkC* | intergenic |
| *S82-S83-glpT-glpQ* | 5’ UTR, riboswitch |

**Table S1. TUs with Mfd binding and increased RpoB occupancy in *Δmfd*.** Associated regulatory RNA categories from previously defined work (6).

| TUs with decreased RpoB in *Δmfd* | regulatory RNA category |
|---|---|
| *srfAA-srfAB-comS-srfAC-srfAD* | none |
| *rsbR-rsbS-rsbT-rsbU-rsbV-rsbW-sigB-rsbX* | none |
| *S1343-csbA-S1342* | 5’ UTR, 3’ UTR |
| *Hpf* | none |
| *ytxG-ytxH-ytxJ* | none |
| *ywjC-S1446* | 3’ UTR |
| *mtlA-mtlF-mtlD* | none |
| *S408-yjbC-S409-spx* | 5’ UTR, intergenic |
| *S928-mgsR* | 5’ UTR |
| *S426-yjcD (asS427-yjzE)* | asRNA |
| *Ctc* | none |
| *ywiE-ywjA-ywjB* | none |
| *S294-csbB* | 5’ UTR |
| *ypiA-ypiB* | none |
| *ahpF-ahpC* | none |
| *ybeC* | none |
| *sunA-sunT-bdbA-yolJ-bdbB* | none |
| *Ybyb* | none |
| *ptsG-ptsH-ptsI* | none |
| *pdaC* | none |
| *gtaB-S1363* | 3’ UTR |
| *S476-ykoM* | 5’ UTR |
| *serA* | none |
| *rsbRD (as927)* | asRNA |
| *S1366* | independent transcript |
| *yjbB* | none |
| *Icd* | none |
| *S1171-ytkA-S1172-dps* | 5’ UTR, intergenic |
| *yjcH-yjcG-yjcF* | none |
| *S1301* | independent transcript |
| *ykzB-ykoL* | none |

**Table S2. TUs with decreased RpoB association in *Δmfd*.** Associated regulatory RNA categories from previously defined work (6).

| *B. subtilis* TA gene | Mfd ChIP-seq association | *Δmfd* RpoB ChIP-seq fold enrichment |
|---|---|---|
| *txpA* (toxin) | **18.1091** | **3.6902** |
| RatA (antitoxin) | **13.3760** | **3.4526** |
| *bsrG* (toxin) | **3.1852** | **2.7718** |
| SR4 (antitoxin) | **4.4111** | **2.8398** |
| *bsrE* (toxin) | 1.9520 | 1.4331 |
| SR5 (antitoxin) | 2.1053 | 1.2556 |
| *yonT* (toxin) | 1.8159 | **3.6664** |
| as-yonT (antitoxin) | 1.7007 | **3.7755** |
| *bsrH* (toxin) | **10.4646** | **3.5415** |
| *as*-bsrH (antitoxin) | **11.4629** | **3.2391** |

**Table S3. Mfd association and RpoB occupancy of *Δmfd* strains at toxin-antitoxin genes.** Genes with bolded values fulfill criteria for significant differences in Mfd occupancy (defined as genes with an Mfd ChIP association one standard deviation greater than the mean) and/or significant increase in RpoB occupancy in *Δmfd* (criteria defined in detail in dataset S2).

| Strain | Genotype and Features | Reference |
|---|---|---|
| HM1 | WT *B. subtilis* JH642 | Brehm et al. J Bacteriol.1973 (7) |
| HM712 | *B. subtilis* 168 Δ*greA*::mls | Koo et al Cell Syst. 2017 (Bacillus Genetic Stock Center) (8) |
| HM1333 | *E. coli* K-12 Δ*mfd*::kan | Baba et al. Mol Syst Biol. 2006 (Coli Genetic Stock Center) (9) |
| HM1451 | *E. coli* MG1655 | Blattner et al. Science. 1997 (10) |
| HM2295 | *E. coli* F' (Kan) placOL2–62-lacZ | Dove et al Nature. 1997 (2) |
| HM2521 | *B. subtilis* JH642 Δ*mfd*::mls | Ragheb et al Mol Cell. 2019 (4) |
| HM2602 | *E. coli* F' (Kan) placOL2–62-lacZ pSIM27(tet) | Ragheb et al Mol Cell. 2019 (4) |
| HM2747 | *E. coli* F' (Kan) placOL2–62-Nanoluc(hyg) | Ragheb et al Mol Cell. 2019 (4) |
| HM2769 | *E. coli* F' (Kan) placOL2–62-Nanoluc(hyg) pSIM27(tet) pHM430(cm)pHM439(amp) | This study |
| HM2771 | *E. coli* F' (Kan) placOL2–62-Nanoluc(hyg) pSIM27(tet) pHM430(cm) pBRα (amp) | This study |
| HM2773 | *E. coli* F' (Kan) placOL2–62-Nanoluc(hyg) pSIM27(tet) pACλCI (cm) pHM439(amp) | This study |
| HM2916 | *E. coli* AG1111 pminiMAD2-mfdL522A | This study |
| HM2932 | *B. subtilis* JH642 MfdL522A | This study |
| HM2965 | *E. coli* F' (Kan) placOL2–62-Nanoluc(hyg) pSIM27(tet) pHM431(cm) pHM439(amp) | This study |
| HM3157 | *B. subtilis* JH642 Mfd-1xmyc | This study |
| HM3808 | *B. subtilis* JH642 Δ*greA*::mls | This study |
| HM3933 | *E. coli* MG1655 Δ*mfd* | This study |
| HM3947 | *B. subtilis* JH642 MfdL522A-1xmyc | This study |

| | | |
|---|---|---|
| HM3986 | *E. coli* DH5α pHM676 | This study |
| HM3988 | *B. subtilis* JH642 amyE::$P_{spank}$-txpA | This study |
| HM3990 | *B. subtilis* JH642 amyE::$P_{spank}$-txpA *Δmfd::mls* | This study |
| HM4002 | *E. coli* DH5α pHM682 | This study |
| HM4003 | *B. subtilis* JH642 amyE::$P_{spank}$-bsrH | This study |
| HM4004 | *B. subtilis* JH642 amyE::$P_{spank}$-bsrH *Δmfd::mls* | This study |
| **Plasmids** | **Description** | **Reference** |
| pBRα | Used as a negative control in bacterial 2-hybrid assays | Dove et al Nature. 1997 (2)(Addgene 53731) |
| pBRα-β-flap | Used to clone and express RNA polymerase α-subunit fusions in *E. coli* | Dove et al Nature. 1997 (2)(Addgene 53734) |
| pCAL215 | Used to amplify erm cassette | Auchtung et al Mol Micro. 2007 (11) |
| pACλCI | Used as a negative control in bacterial 2-hybrid assays | Dove et al Nature. 1997 (2)(Addgene 53730) |
| pACλCI-β-flap | Used to clone and express λCI fusions in *E. coli* | Dove et al Nature. 1997 (2)(Addgene 53733) |
| pHM430 | Plac-CI-Bsubmfd(494-625) | This study |
| pHM431 | Plac-CI-Bsubmfd(494-625)L522A | This study |
| pHM439 | Plac-a-BsubrpoB(21-131) | This study |
| pHM676 | $amp^R$, amyE::$P_{spank(hy)}$-txpA, lacI, $spec^R$ | This study |
| pHM682 | $amp^R$, amyE::$P_{spank(hy)}$-bsrH, lacI, $spec^R$ | This study |
| pHM707 | pminiMAD2-BsubMfdL522A | This study |
| pDR111 | $amp^R$, amyE::$P_{spank(hy),}$ lacI, $spec^R$ | Guérout-Fleury et al Gene. 1996 (12) |
| pminiMAD2 | Scarless integration plasmid for *B. subtilis* | Patrick and Kearns Mol Micro. 2008 (1) |
| pNL1.1 | NanoLuc expression vector | Promega (GenBank Accession #JQ513379) |

**Table S4. Bacterial strains and plasmids used in this study.**

| Primer # | Sequence | Description |
|---|---|---|
| HM80 | AGGATAGGGTAAGCGCGGTATT | *B. subtilis* rRNA qPCR |
| HM81 | TTCTCTCGATCACCTTAGGATTC | *B. subtilis* rRNA qPCR |
| HM192 | CCGTCTGACCCGATCTTTTA | *B. subtilis yhaX* qPCR |
| HM193 | GTCATGCTGAATGTCGTGCT | *B. subtilis yhaX* qPCR |
| HM910 | AAGGCACATGGCTGAATATCG | *B. subtilis lacZ* qPCR |
| HM911 | ACACCAGACCAACTGGTAATGG | *B. subtilis lacZ* qPCR |
| HM1004 | CATGAGGGTACCGATGATCAGCGGT CAATTGA | For amplifying *B. subtilis mfd* to insert into pminiMAD2 |
| HM1005 | CATGAG GGATCCCATAGTGCTGCT GTGCCAA | For amplifying *B. subtilis mfd* to insert into pminiMAD2 |
| HM1555 | CAGGTCAACTAGTTCAGTATGGACGACAC | *B. subtilis rpoB* qPCR |
| HM1556 | CTCTAAGACCCTCATCAAGAAACCACTG | *B. subtilis rpoB* qPCR |
| HM3286 | AGTGGCCTGAAGAGACGTTTGGCGCA AAAAGCTATTCTGAGCTTCAAATTG | For amplifying *B. subtilis mfd* (bp 1483-1875) with homology to pACCI for Gibson and extra base to maintain frame. |
| HM3287 | CTGCGATGCAGATCTGTAAGGTAAGTT AAGTCTCTTGATAAGGGAAAGCC | For amplifying *B. subtilis mfd* (bp 1483-1875) with homology to pACCI for Gibson with stop codon added |
| HM3292 | AAGTGAAAGAAGAGAAACCAGAGGCA GAAGTGTTAGAATTACCAAATCTCATT G | For amplifying *B. subtilis rpoB* (bp 64-393) with homology to pBRa for Gibson and extra base to maintain frame. |
| HM3293 | CGGCCACGATGCGTCCGGCGTAGAGT TATTCCGCACCGTTAATGATAAAAG | For amplifying *B. subtilis rpoB* (bp 64-393) R with homology to pBRa for Gibson and stop codon added. |
| HM3540 | GAATGCCGTTGATTTCAGCAGTTTCAA TCCCCAGGTATTTTCCG | Quickchange primer to make L522A *mfd* mutation |
| HM3541 | CGGAAAATACCTGGGGATTGAAACTG CTGAAATCAACGGCATTC | Reverse quickchange primer to make L522A *mfd* mutation |
| HM3759 | CAAGTCCTCTTCACTGATTAACTTCTG CTCCGTTGATGAAATGGTTTGCT | For amplifying C-terminally myc tagged *mfd* by SOE PCR |
| HM3760 | GAGCAGAAGTTAATCAGTGAAGAGGA CTTGTAAATTTTGTTACTCTCTGGTGTA TATTAC | For amplifying C-terminally myc tagged *mfd* by SOE PCR |
| HM3854 | CGAGGCTCCTGTCACTGCT | For amplifying erm-HI cassette |
| HM3969 | GAGCAGAAGTTAATCAGTGAAGAGGA CTTGATTTTGTTACGCAGGCGAGAAAG GAGAGAG | For amplifying erm-HI cassette with myc tag at 5' end |

| | | |
|---|---|---|
| HM5162 | ACACTCCTCATGTTTGCCTT | *B. subtilis tnrY* qPCR |
| HM5163 | GTGTCGGCGGTTCGATT | *B. subtilis tnrY* qPCR |
| HM5418 | CATGATGCTAGCTGAAAGGAGGTGAAATTATGTCGAC | For making *txpA* overexpression construct cloning |
| HM5419 | CATGATGCATGCCTACCCTTTAATAGGAGGGT | For making *txpA* overexpression construct cloning |
| HM5437 | CAAGCAAAAGTATTGCAACT | *B. subtilis ratA* qPCR |
| HM5438 | GGTAATGTGGTAATGTGGTA | *B. subtilis ratA* qPCR |
| HM5441 | ATGTCGACCT ATGAATCTCT | *B. subtilis txpA* qPCR |
| HM5442 | CCCATGTCATAATCCCGCCT | *B. subtilis txpA* qPCR |
| HM5443 | TTACTGTAAAGGAAAAGTGT | *B. subtilis txpA* qPCR |
| HM5444 | CTACCCTTTAATAGGAGGGT | *B. subtilis txpA* qPCR |
| HM5462 | CATGATGCTAGCATGGTTTAGTATAAATGAAT | For making *bsrH* overexpression construct cloning |
| HM5463 | CATGATGCATGCAAGAGACCCGGTTGCCGCCGGG | For making *bsrH* overexpression construct cloning |
| HM5156 | ATAATGATGATTGTAACGTCAAGCC | *B. subtilis yolA* qPCR |
| HM5157 | GCCTAACCCTTCAGGTGTC | *B. subtilis yolA* qPCR |
| HM5571 | CCGCCGGGTCAGTATAAATG | *B. subtilis bsrH* qPCR |
| HM5572 | CCCTTGAGCTCGGCAAAG | *B. subtilis bsrH* qPCR |

**Table S5. Oligonucleotides used in this study.**

**Dataset S1 (separate file)** Quantification of Mfd association of genes in the *B. subtilis* 168 genome. Mfd-myc binding was calculated by taking the average read count across a given gene and normalizing internally to overall read counts as well as to WT *B. subtilis* (lacking a myc tag). Values were subsequently $\log_2$ normalized. Genes are sorted from highest to lowest Mfd binding values. Those genes with greater than one standard deviation from the mean Mfd-myc binding value were defined as Mfd associated.

**Dataset S2 (separate file)** Genes with increased RpoB ChIP association in *B. subtilis* Δ*mfd*, sorted by increasing p-value. (logFC= log-fold change, logCPM= log counts per million, FDR= false discovery rate). The following criteria were used to define increased RpoB association= logFC>1, logCPM>4, p-value< $1x10^{-4}$, FDR< .001. Genes in bold text are also Mfd associated.

**Dataset S3 (separate file)** Genes with decreased RpoB ChIP association in *B. subtilis* Δ*mfd*, sorted by increasing p-value. Criteria used to define decreased RpoB association is the same as described in Dataset S2.

**Dataset S4 (separate file)** Genes with increased RpoB ChIP association in *E. coli* Δ*mfd*, sorted by increasing p-value. (logFC= log-fold change, logCPM= log counts per million, FDR= false discovery rate). The following criteria were used to define increased RpoB association= logFC>1, logCPM>3, p-value< $1x10^{-4}$, FDR< .001. Genes in bold text are also Mfd associated.

**Dataset S5 (separate file)** Genes with decreased RpoB ChIP association in *E. coli* Δ*mfd*, sorted by increasing p-value. Criteria used to define decreased RpoB association is the same as described in Dataset S4. Genes in bold text are also Mfd associated.

**Dataset S6 (separate file)** Genes with altered RpoB ChIP association in Δ*greA*. Genes are sorted by increasing p-value, with the first 12 genes exhibiting increased RpoB occupancy in Δ*greA* and the remaining genes exhibiting decreased RpoB occupancy in Δ*greA*. To define significant differences, the same criteria were used as described in Tables S1 and S2.

**Dataset S7 (separate file)** Upregulated and Downregulated genes in *B. subtilis Δmfd* strain based on DEseq2 analysis. Genes are sorted by increasing p-value. The following criteria was used to define transcriptional differences = logFC> 1, logCPM> 2, FDR< .05
